# Comparative analysis, applications, and interpretation of electronic health record-based stroke phenotyping methods

**DOI:** 10.1101/565671

**Authors:** Phyllis M. Thangaraj, Benjamin R. Kummer, Tal Lorberbaum, Mitchell V. S. Elkind, Nicholas P. Tatonetti

**Affiliations:** Department of Biomedical Informatics, Columbia University, New York, NY; Department of Systems Biology, Columbia University, New York, NY; Department of Neurology, Icahn School of Medicine at Mt. Sinai, New York, NY; Department of Neurology, Vagelos College of Physicians and Surgeons, Columbia University, New York, NY; Department of Epidemiology, Mailman School of Public Health, Columbia University, New York, NY

**Keywords:** phenotyping algorithms, acute ischemic stroke, machine learning, electronic health record studies

## Abstract

**Background and Purpose:** Accurate identification of acute ischemic stroke (AIS) patient cohorts is essential for a wide range of clinical investigations. Automated phenotyping methods that leverage electronic health records (EHRs) represent a fundamentally new approach cohort identification. Unfortunately, the current generation of these algorithms is laborious to develop, poorly generalize between institutions, and rely on incomplete information. We systematically compared and evaluated the ability of several machine learning algorithms and case-control combinations to phenotype acute ischemic stroke patients using data from an EHR.

**Methods:** Using structured patient data from the EHR at a tertiary-care hospital system, we built machine learning models to identify patients with AIS based on 75 different case-control and classifier combinations. We then determined the models’ classification ability for AIS on an internal validation set, and estimated the prevalence of AIS patients across the EHR. Finally, we externally validated the ability of the models to detect self-reported AIS patients without AIS diagnosis codes using the UK Biobank.

**Results:** Across all models, we found that the mean area under the receiver operating curve for detecting AIS was 0.963±0.0520 and average precision score 0.790±0.196 with minimal feature processing. Logistic regression classifiers with L1 penalty gave the best performance. Classifiers trained with cases with AIS diagnosis codes and controls with no cerebrovascular disease diagnosis codes had the best average F1 score (0.832±0.0383). In the external validation, we found that the top probabilities from a model-predicted AIS cohort were significantly enriched for self-reported AIS patients without AIS diagnosis codes (65-250 fold over expected).

**Conclusions:** Our findings support machine learning algorithms as a way to accurately identify AIS patients without relying on diagnosis codes or using process-intensive manual feature curation. When a set of AIS patients is unavailable, diagnosis codes may be used to train classifier models. Our approach is potentially generalizable to other academic institutions and further external validation is needed.

## INTRODUCTION

Stroke is a complex disease that is a leading cause of death and severe disability for millions of survivors worldwide.^1^ Accurate identification of stroke etiology, which is most commonly ischemic but encompasses several other causative mechanisms, is essential for risk stratification, optimal treatment, and support of clinical research. While electronic health records (EHR) are an emerging resource that can be used to study stroke patients, identification of stroke patient cohorts using the EHR requires the integration of multiple facets of data, including medical notes, labs, imaging reports, and medical expertise of neurologists. This process is often manually performed and time-consuming, and can reveal mis-classification errors.^2^ One simple approach to identify acute ischemic stroke (AIS) is the diagnosis-code based algorithm created by Tirschwell and Longstreth.^3^ However, identifying every AIS patient using these criteria can be difficult due to the inaccuracy and incompleteness of diagnosis recording through insurance billing. ^3–5^ Additionally, this approach prevents the identification of AIS patients until after hospital discharge, thereby limiting the clinical usability of identification algorithms in time-sensitive situations, such as in-hospital care management, research protocol enrollment, or acute treatment.

Reproducibility and computability of phenotyping algorithms stem from the use of structured data, standardized terminologies, and rule-based logic.^6^ Phenotyping features from the EHR have been traditionally culled and curated by experts to manually construct algorithms, ^7^ but machine learning techniques present the potential advantage of automating this process of feature selection and refinement.^8,9^ Recent machine learning approaches have also combined publicly available knowledge sources with EHR data to facilitate feature curation.^10,11^ Additionally, while case and control phenotyping using EHR data has also relied on a small number of expert curated cohorts, recent studies have demonstrated that ML approaches can identify such cohorts using automated feature selection and imperfect case definitions in a high-throughput manner.^12–14^ Two stroke phenotyping algorithms have also used machine learning to enhance the classification performance of a diagnosis-code based AIS phenotyping algorithm.^15,16^ However, while ML models present an opportunity to automate identification of AIS patients (i.e. phenotyping) with commonly accessible EHR data and develop new approaches to etiologic identification and subtyping, the optimal combination of cases and controls to train such models remains unclear.

Given the limitations of manual and diagnosis-code cohort identification, we sought to develop phenotypic classifiers for AIS using machine learning approaches, with the objective of specifically identifying AIS patients that were missing diagnosis codes. Additionally, considering the challenge of identifying true controls in the EHR for the purpose of model training, we also attempted to determine the optimal grouping of cases and controls by selecting and comparing model discriminatory performance with multiple case-control group combinations. We also sought to contrast model training based on cases defined by diagnostic code with that using manually-curated cohorts. Our phenotyping method utilizes machine learning classifiers with minimal data processing to increase the number of stroke patients recovered within the EHR and reduce the time and effort needed to find them for research studies.

## METHODS

### Study Design

In this study, we developed several machine learning phenotyping models for AIS using combinations of different case and control groups derived from our institution’s EHR data. Use of patient data was approved by Columbia’s institutional review board. We also applied key methods to optimize number of features for generalizability, as well as calibration to ensure a clinically meaningful model output, and model robustness to missing data. To estimate the prevalence of potential AIS patients without AIS-related *International Classification of Diseases* (ICD) codes, we then applied the developed models to all patients in our institutional EHR. Finally, we externally validated our best-performing model in an independent cohort from the UK Biobank to evaluate its ability to detect self-reported AIS patients without the requisite ICD codes. Figure 1 shows the overall workflow of training and testing the models, the models’ evaluation, and its testing in an independent test set.

**Figure 1:**
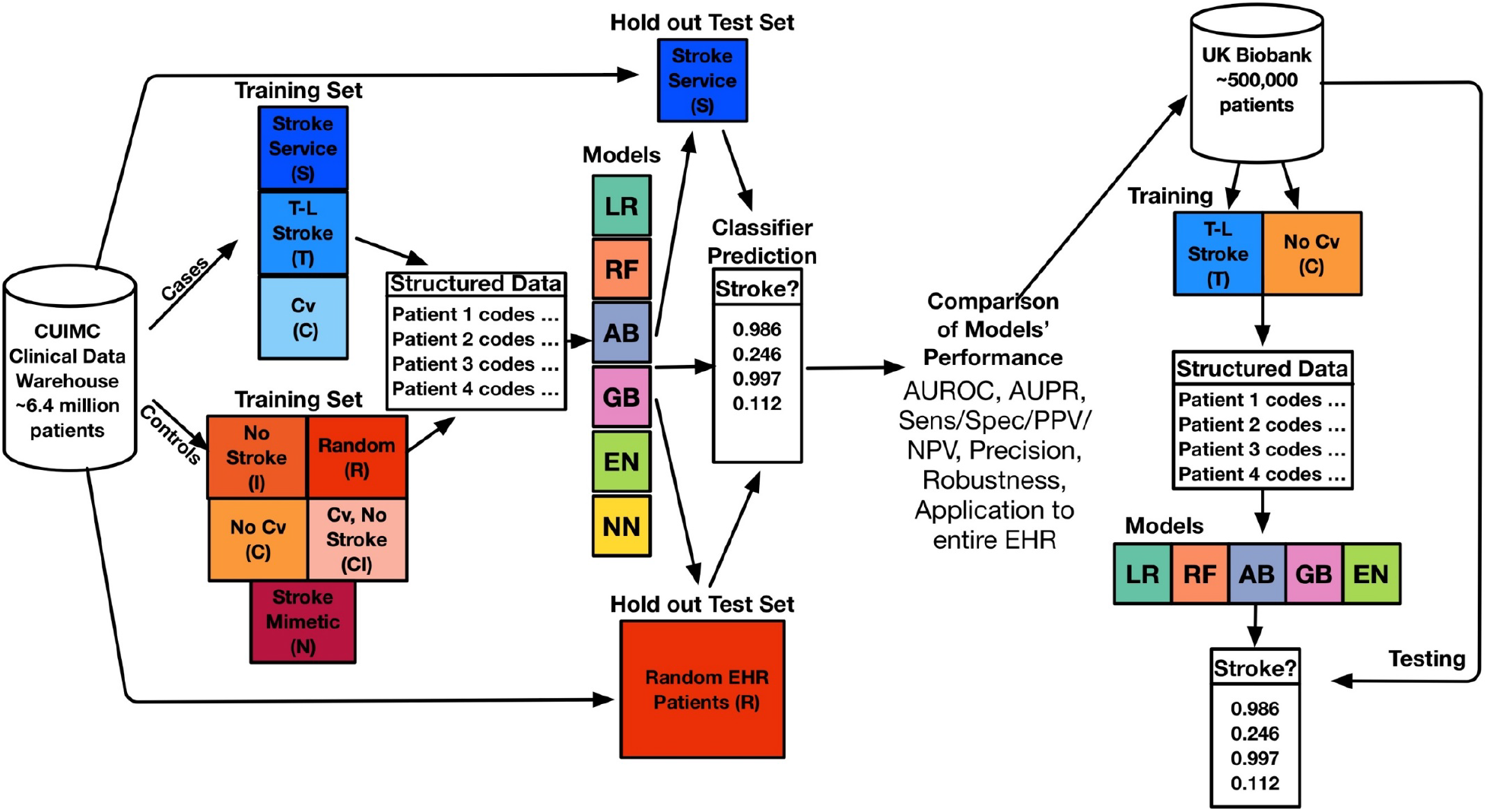
Schematic of Model Training, Testing, Evaluation, and Application to UK Biobank. See methods for case/control abbreviations. Case: Control ratio was 1:1 and models included Random Forest (RF), Logistic Regression with L1 penalty (LR), Neural Network (NN), Gradient Boosting (GB), Logistic Regression with Elastic Net Penalty (EN) and Adaboost (AB). AUROC: Area Under the Receiver Operating Curve, AUPR: Area under the Precision-Recall Curve, Sens: Sensitivity, Spec: Specificity, PPV: Positive Predictive Value, NPV: Negative Predictive Value.

### Data Sources

We used data from patients in the Columbia University Irving Medical Center Clinical Data Warehouse (CUIMC CDW), which contains longitudinal health records of 6.4 million patients from CUIMC’s EHR, spanning 1985-2018. This includes patients from the CUIMC stroke service (Figure 1, Table 1), that were part of a larger group of patients with acute cerebrovascular diseases and were prospectively identified and recorded as part of daily research activities by a CUIMC stroke physician between 2011 and 2018. Two researchers (PT and BK) each manually reviewed 50 patients’ charts from this cohort to determine baseline false positive rates.

**Table 1:**
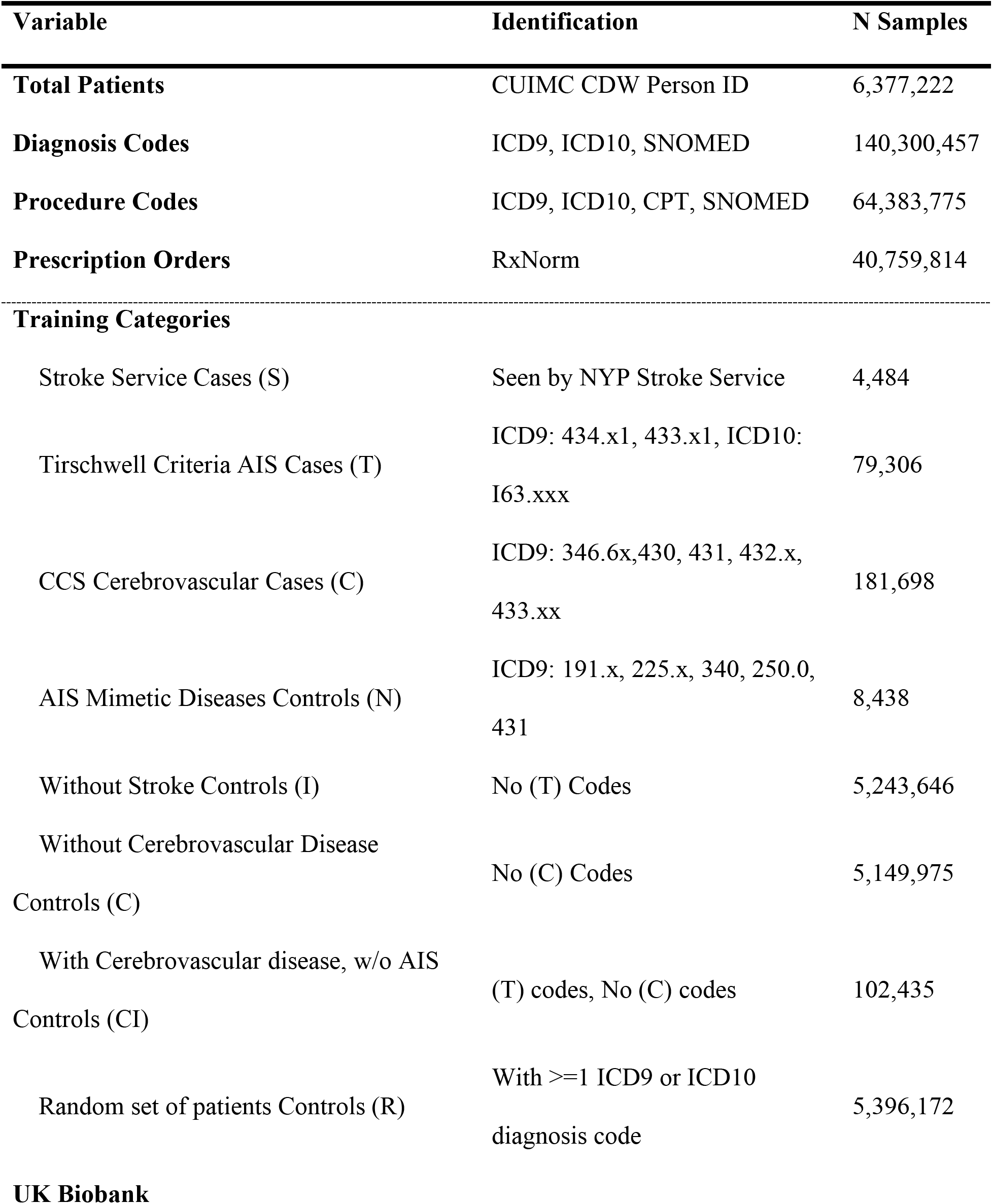
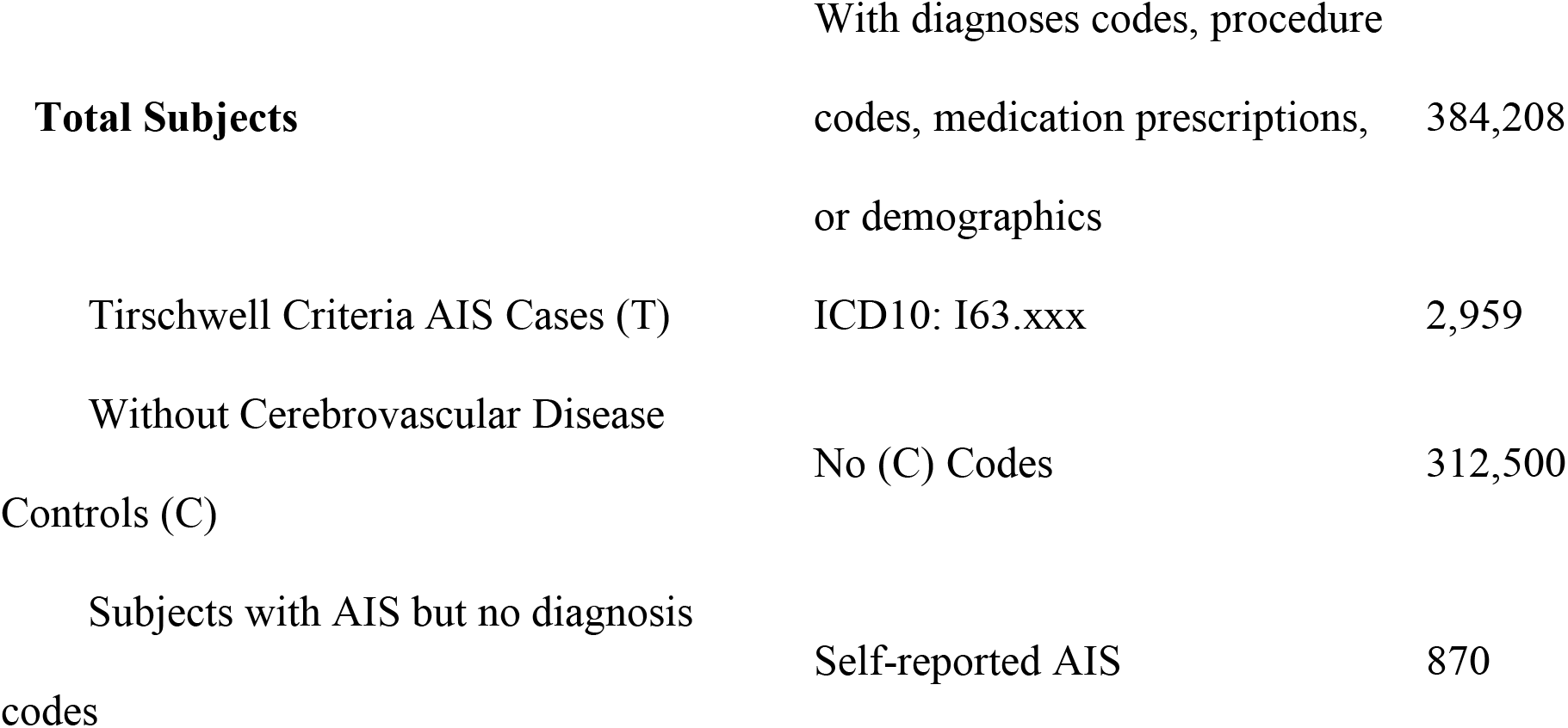
Select Structured Data and Sample Case/Controls for models available in CUIMC Common Data Warehouse.

### Patient Population

We defined 3 case groups. We first included all patients from the CUIMC stroke service that were recorded as having AIS (cohort S). We then defined all patients in the CDW that met the Tirschwell-Longstreth (T-L) diagnosis code criteria for AIS (cohort T), which comprise ICD9CM codes 434.x1, 433.x1, 436 (where x is any number) and the code is in the primary diagnostic position. ^3^ Our dataset did not specify the diagnostic position of codes. We also included ICD10 code equivalents, I63.xxx or I67.89, with the ICD10 codes being determined from ICD9 from Centers for Medicaid and Medicare Services (CMS) General Equivalence Mappings.^18^ Because patients with cerebrovascular disease are also likely to have suffered AIS, but may not have an attached AIS-related diagnosis code, we also created a group of cases according to cerebrovascular disease-related ICD codes defined by the *ICD-9-Clinical Modification* (CM) Clinical Classifications Software tool (CCS), as well as their ICD10 equivalents (cohort C).^17^

We then defined 4 control groups (Figure 1, Table 1). First, we defined a control group of patients without AIS-related diagnosis codes (I). Due to the fact that cerebrovascular disease is a major risk factor for stroke,^19^ and to test a more stringent control definition than that of group (I), we also defined an additional group without any of the CCS cerebrovascular disease codes defined in cohort (C). Then, we defined a control set using CCS cerebrovascular disease diagnosis codes other than AIS (CI). Because multiple clinical entities can present as AIS, we also defined a group of controls according to diagnosis codes for AIS mimetic diseases (N), including hemiplegic migraine (ICD9-CM 346.3), brain tumor (191.xx, 225.0), multiple sclerosis (340), cerebral hemorrhage (431), and hypoglycemia with coma (251.0). Finally, we identified a control group culled from a random sample of patients (R).

### Model Features

From the CDW, we gathered race, ethnicity, age, sex, diagnostic and procedure insurance billing codes as well as medication prescriptions for all patients. We dichotomized each feature based on its presence or absence in the data. Because *Systematized Nomenclature of Medicine* (SNOMED) concept IDs perform similarly to ICD9 and ICD10 codes for phenotyping, ^20^ we mapped diagnoses and procedure features from ICD9, ICD10, and *Current Procedural Terminology 4* (CPT4) codes to SNOMED concept IDs, and used *RxNorm* IDs for medication prescriptions. We identified patients with Hispanic ethnicity using an algorithm combining race and ethnicity codes.^21^ The most recent diagnosis in the medical record served as the age end point and we dichotomized age as greater than or equal, or less than 50 years. We excluded from our feature set any diagnosis codes that were used in any case or control definitions. Because approximately 5 million patients exist in the CUIMC CDW without a cerebrovascular disease diagnosis code, we addressed this large resultant imbalance in cases and controls by randomly sampling controls to create a balanced, or 1:1 case to control ratio. In addition, we set the maximum sample size to 16,000 patients in order to control the size of the feature set.

#### Model Development

Using all 15 case-control combinations, we trained 75 models using logistic regression classifiers with L1 and elastic net regularization, as well as random forest, AdaBoost, gradient boosting, and neural network classifiers on the gathered features. We chose these classifiers to compare a variety of feature-to-outcome relationships: linear (logistic regression), ensemble (random forest, AdaBoost, gradient boosting), and non-linear (neural network). We tuned the models’ hyperparameters using 10-fold cross validation (Supplementary Methods). We then determined a probability threshold for each model using the training set. Within the validation set of each training fold, controls were bootstrapped to form a 100:1 control to case ratio to represent the prevalence of AIS in the general population.^22^ Precision and recall were then calculated from the bootstrapped set. To determine the optimal threshold to maximize precision and recall, we calculated maximum F scores at different *β*s using the following equation

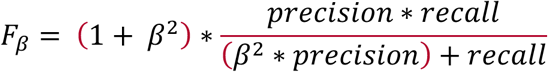

where ß takes the values 1.0, 0.5, 0.25, and 0.125 with increasing weight of precision. Using the probability threshold determined from cross-fold validation, we then calculated the maximum F1 score, sensitivity, specificity, positive predictive value, negative predictive value, and precision on a holdout set of 1000 patients from the stroke service and 100,000 non-overlapping randomly selected patients. We chose this test ratio to imitate the prevalence of AIS in the general population. The models were evaluated on the test set with area under the receiver operating curve (AUROC) and average precision score (AP), a proxy for area under the precision-recall curve. All models were programmed using the Python sklearn scientific computing package (Python Software Foundation, www.python.org).^23^ We then aggregated common features found in the top ten in importance or beta coefficient weight for each model, and we evaluated the contribution of each feature to each model by comparing its prevalence in the cases with its prevalence in the controls and as a function of its importance (or weight) in the model.

### Internal Validation using All EHR Patients

To identify the number of patients classified as having AIS in our institutional EHR, we applied each of the 75 models to the entire patient population in the CUIMC CDW with at least one diagnosis code. We chose a probability threshold based on the maximum F1 score determined for each model from the training set. We also determined the percentage of patients that had AIS ICD9 codes as defined by T-L criteria and associated ICD10 codes.

### External Validation

The UK Biobank is a prospective health study of over 500,000 participants, ages 40-69, containing comprehensive EHR and genetic data.^24^ Given that this dataset contains 2,959 patients with an AIS related ICD10 code, similar to our T case cohort, and 870 patients with self-reported AIS, without AIS related ICD10 codes, the UK Biobank is an ideal cohort to evaluate our machine learning models’ ability to recover potential AIS patients that lack AIS-related ICD10 codes. We chose the most accurate and robust case-control combination from our models (cases defined by the T-L AIS codes (T) and controls without codes for cerebrovascular disease (C) in a 1:1 case-control ratio as our training set) to train the phenotyping model using conditions specified by ICD10 codes, procedures specified by OCPS4 codes, medications specified by RxNorm codes, and demographics as features, excluding self-reported features as well as those that were used to create the cohorts. We then tested our models on the rest of the UK Biobank data which included self-reported AIS cases. We resampled the control set 50 times and evaluated the performance of the algorithm through AUROC, AP, and precision at the top 50, 100, 500 and 870 patients (ordered by model probability).

## RESULTS

### Study Cohort

Table 1 presents the data and the total number of patients available for each set of cases and controls used in the training and internal and external validation parts of this study. Out of the CUIMC EHR, which has a total of 6.4 million patients, we extracted 4,844 stroke service patients, which we found to have a 4-16% false positive rate for stroke. Supplementary Table 2 presents demographic characteristics.

### Algorithm Performance

We trained 75 models using all combinations of cases, controls, and model types after excluding 15 neural network models due to poor performance. Logistic regression classifiers with L1 penalty gave the best AUROC performance (0.913-0.997) and the best average precision score (0.662-0.969), followed by logistic regression classifiers with elastic net penalty (Figure 2, Supplementary Table 3).

**Figure 2:**
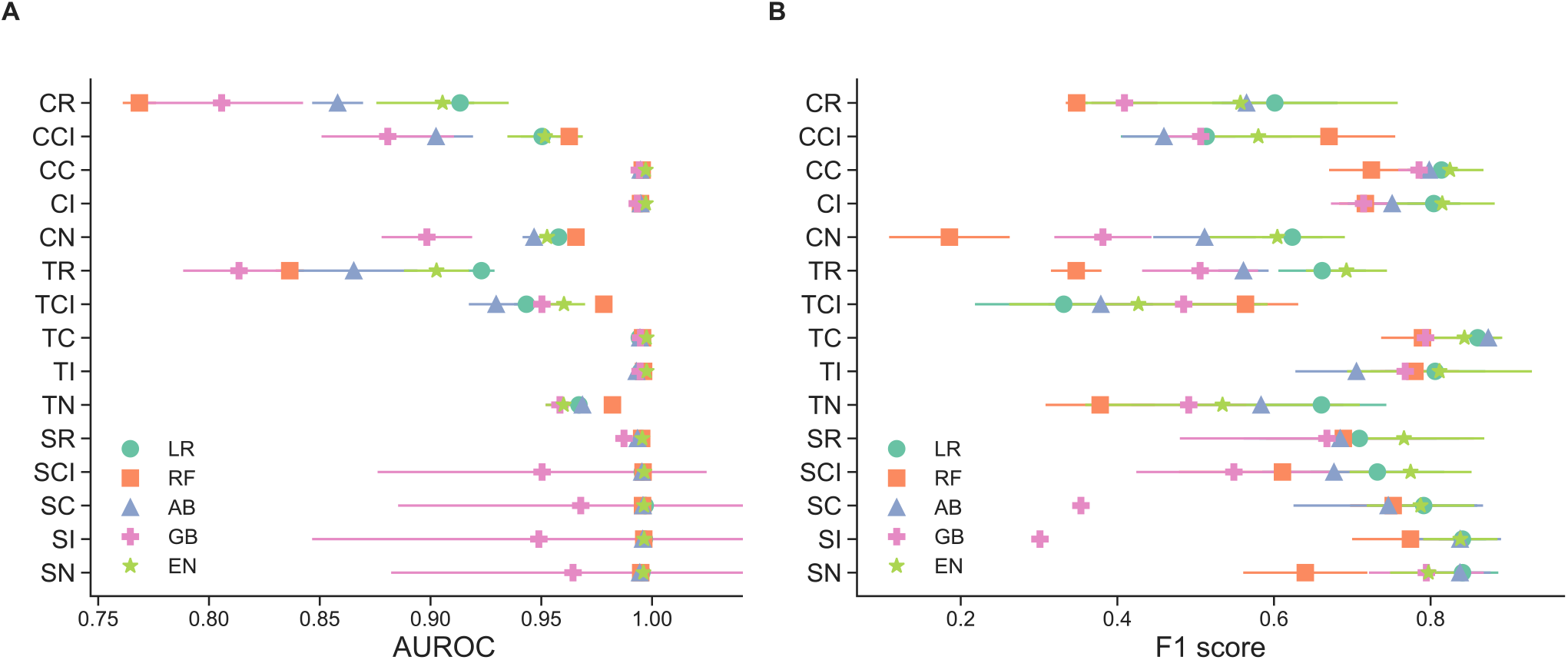
Performance of select models on holdout test set ((a): AUROC, (b): F1). Different combinations of cases and controls are shown on the y-axis. (LR) logistic regression with l1 penalty, (RF) random forest, (AB) AdaBoost, (GB) gradient boosting, (EN) logistic regression with elastic net penalty. Different combinations of cases and controls are shown on the y-axis. Cases (first letter) may be one of cerebrovascular (C), T-L (T), or Stroke Service (S). Controls (second and third letters) may be one of random (R), cerebrovascular disease but no AIS code (CI), no cerebrovascular disease (C), no AIS code (I), or a stroke mimetic disease (N), See Methods and Supplementary Table 1 for definitions of sets. Threshold to compute the F1 score on the testing set was chosen as the threshold that yielded the maximum F1 in cross-validation on the training set (Methods, Supplementary Table 6).

Across all classifier types, the models using the T-C case-control combination had the best average F1 score (0.832±0.0383), whereas logistic regression models with L1 penalty (LR) and elastic-net penalty had the best classifier average F1 score (0.705±0.146 and 0.710±0.134 respectively) (Figure 2B, Supplementary Table 6). Use of cases from the CUIMC stroke service gave the highest average precision (0.932±.0536), while cases identified through AIS diagnosis codes and controls without cerebrovascular disease or AIS-related diagnosis codes (TC, TI) gave high precision as well (0.896±0.0488 and 0.918±0.0316, respectively). The sensitivity of the models ranged widely, between 0.18 and 0.96, while specificity narrowly ranged between 0.993-1.0 (Supplementary Table 7).

### Feature Importance

We found the most commonly chosen features associated with stroke diagnosis were procedures used in evaluation of AIS, including extra- and intra-cranial arterial scans, CT scans and MRIs of the brain, and MR angiography (Figure 3A). We also found that all 75 models relied on incremental contributions from many different features (Figure 3B, Supplementary Figures 20-34).

**Figure 3:**
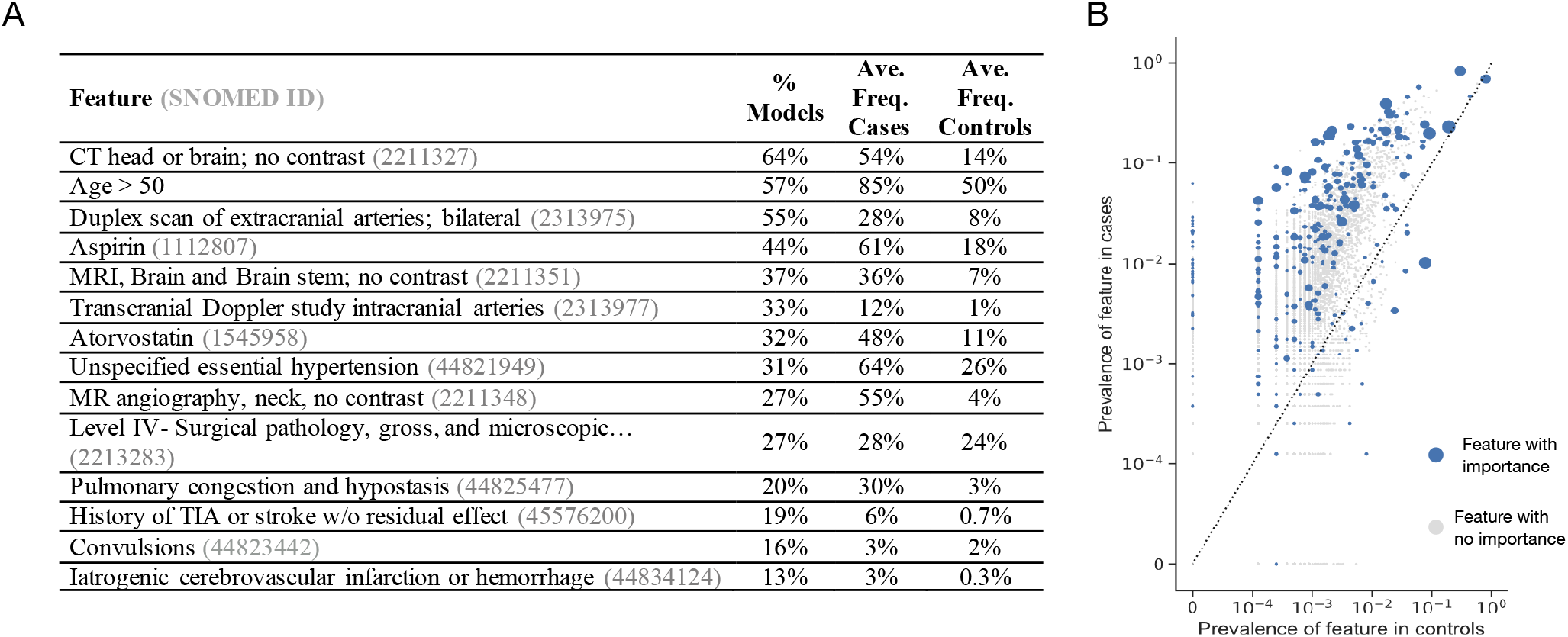
**A:** Common top 10 features in the models. After each of the 75 models were trained, we counted the number of times each feature was represented as one of the top ten by absolute coefficient weight, for methods like logistic regression, or by feature importance, for methods like random forest. Above are features from this analysis along with the proportion of models in which they were in the top ten (% Models), the average frequency in the cases (Ave. Freq. Cases) and the average frequency in the controls (Ave. Freq. Controls). **B:** Prevalence of features in cases vs controls in the TC AB model. Axes were on a logarithmic scale. Increasing size of blue dot correlates with higher feature importance or beta coefficient weight, depending on the classifier type. Gray dots are features with zero importance.

### Internal Validation in Institutional EHR

We applied the 75 models to the entire CUIMC EHR with at least one diagnosis code, totaling between 5,324,725 and 5,315,923 patients depending on the case/control set. We found that the results varied widely across models, but most predicted a prevalence of between 0.2-2% of patients in the EHR were AIS patients. The models with controls with cerebrovascular disease codes but no AIS codes predicted the lowest prevalence of AIS patients, and found 50.3-100% of the proposed patients had AIS diagnosis codes. The models with the best performance and robustness, 1) stroke service cases and controls without cerebrovascular disease codes and 2) cases with AIS codes and controls without cerebrovascular disease codes with 1) Logistic Regression and L1 Penalty classifier and 2) Adaboost classifier, had sensitivities between 0.822-0.959, specificities 0.994-0.999, and estimated AIS prevalence in the EHR ranging between 1.3-2.0% (Supplementary Table 7, Table 2). Within these proposed AIS patients, 37.7-41.4% had an AIS diagnosis code (Table 2).

**Table 2.** Prevalence of acute ischemic stroke patients identified by each classifier across the EHR and proportion of those patients with T-L criteria. Prev=prevalence. See Supplementary Table 1 for case-control and model abbreviations’ definitions.

| **Case/** | **LR** | **RF** | **AB** | **GB** | **EN** | **LR** | **RF** | **AB** | **GB** | **EN** |
| --- | --- | --- | --- | --- | --- | --- | --- | --- | --- | --- |
| **Control** | **EHR** | **EHR** | **EHR** | **EHR** | **EHR** | **with** | **with** | **with** | **with** | **with** |
| **Combo** | **Prev** | **Prev** | **Prev** | **Prev** | **Prev** | **AIS** | **AIS** | **AIS** | **AIS** | **AIS** |
|  |  |  |  |  |  | **codes** | **codes** | **codes** | **codes** | **codes** |
| **SN** | 0.7 | 0.7 | 1.0 | 1.3 | 0.7 | 41.3 | 32.2 | 35.6 | 29.0 | 26.4 |
| **SI** | 1.1 | 2.0 | 1.5 | 1.7 | 1.1 | 40.5 | 23.0 | 35.7 | 29.8 | 27.1 |
| **SC** | 1.3 | 1.7 | 1.5 | 1.8 | 1.3 | 37.7 | 25.4 | 37.9 | 30.8 | 28.5 |
| **SCI** | 0.2 | 0.1 | 0.2 | 0.3 | 0.2 | 83.1 | 82.6 | 76.9 | 72.2 | 63.5 |
| **SR** | 0.2 | 0.2 | 0.3 | 0.5 | 0.2 | 75.4 | 63.2 | 68.8 | 58.2 | 48.9 |
| **TN** | 0.9 | 0.8 | 0.9 | 1.0 | 0.9 | 44.7 | 28.5 | 47.2 | 35.6 | 22.5 |
| **TI** | 1.6 | 2.3 | 1.4 | 4.7 | 1.6 | 43.8 | 31.4 | 47.9 | 21.8 | 8.10 |
| **TC** | 1.7 | 2.7 | 2.0 | 1.6 | 1.7 | 41.4 | 28.2 | 39.0 | 43.1 | 32.6 |
| **TCI** | 0.1 | 0.0 | 0.1 | 0.1 | 0.1 | 94.6 | 96.1 | 85.9 | 95.3 | 79.0 |
| **TR** | 0.8 | 0.8 | 0.8 | 0.4 | 0.8 | 46.1 | 40.0 | 44.0 | 61.4 | 31.1 |
| **CN** | 1.3 | 1.3 | 1.3 | 1.0 | 1.3 | 34.0 | 17.1 | 33.5 | 31.5 | 21.4 |
| **CI** | 2.0 | 3.3 | 1.9 | 1.9 | 2.0 | 37.5 | 24.2 | 39.5 | 39.8 | 39.9 |
| **CC** | 2.3 | 3.3 | 2.2 | 2.1 | 2.3 | 35.6 | 25.3 | 37.2 | 37.1 | 29.9 |
| **CCI** | 0.0 | 0.0 | 0.1 | 0.0 | 0.0 | 97.5 | 100 | 50.3 | 92.8 | 74.2 |
| **CR** | 1.0 | 0.9 | 0.9 | 0.7 | 1.0 | 37.3 | 35.6 | 37.7 | 42.6 | 29.6 |

### External Validation

We evaluated the performance of the TC models on identifying 870 patients with self-reported AIS. The top 50, 100, 500, and 870 probabilities had a precision of over 13%, and up to 50% (Figure 4). Since within the test set only 0.2% of the patients had self-reported AIS, this translates to a 60-250-fold increase in AIS detection over random choice.

**Figure 4.**
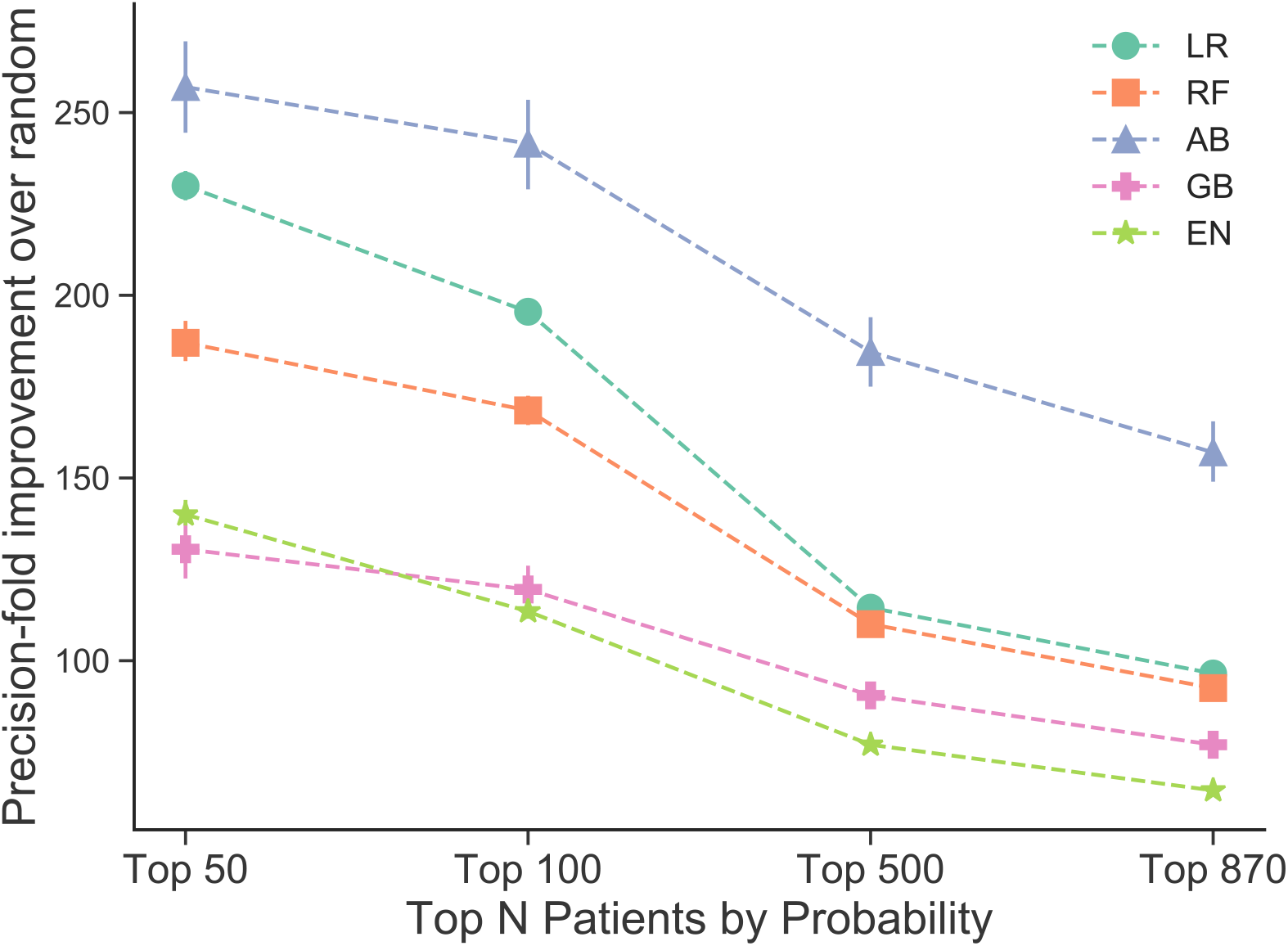
Precision-fold over random sampling of self-reported acute ischemic stroke cases at top 50, 100, 500, and 870 patient probabilities assigned by machine learning algorithms. With 95% confidence intervals in error bars. See Supplementary Table 1 for model abbreviations’ definitions.

## DISCUSSION

Using a feature-agnostic, data-driven approach with minimal data transformation, we developed models that identify acute ischemic stroke (AIS) patients from commonly-accessible EHR data at the time of patient hospitalization without making use of AIS-related ICD9 and ICD10 codes as defined by Tirschwell and Longstreth. In demonstrating that AIS patients can be recovered from other EHR-available structured clinical features without AIS codes, this approach is in contrast to previous machine learning phenotyping algorithms, which have relied on manually curated features or use AIS-related diagnosis codes as the sole nonzero features in their models. ^15,16,3^

Cases and controls for training of phenotyping algorithms can be challenging to identify and define given the richness of available EHR data. From the sparsity of diagnosis codes in the EHR, it follows that patients lacking an AIS-related diagnosis code may not always be considered as a control in stroke cohorts. Similarly, it is difficult to determine whether patients with cerebrovascular diseases, which can serve as risk factors for AIS, or share genetic and pathophysiologic underpinnings with AIS should be considered controls. Additionally, due to the prevalence of AIS mimics, cohort definitions based on diagnosis code criteria may be unreliable. In light of the problems in defining patient cohorts from EHR data, we found marked differences in classifying performance across 15 different case-control training sets. While training with cases from the CUIMC stroke service cases identified stroke patients most accurately and with the highest precision and recall, we also found that training with cases identified from AIS codes with controls from either 1) no cerebrovascular disease or 2) no AIS codes afforded high precision (Supplementary Table 3). These findings suggest that a manually curated cohort may not be necessary to train the phenotyping models, and the AIS codes may be enough to define a training set. Using these models, we also increased our AIS patient cohort by 60% across the EHR, suggesting that the AIS codes themselves are not sufficient to identify all AIS patients.

We found that stroke evaluation procedures, such as a CT scan or MRI, were important features in many of the models. Since none of these models use AIS diagnosis codes as features, this suggests that procedures may serve as proxies for them. In some cases, the AIS code will only be added during outpatient follow up. For example, while in the stroke service set, 13.5% of cases did not have AIS codes in the inpatient setting but did in the outpatient setting, and 90% of these patients had had a CT scan of the head. We also found evidence that procedures provided a significant contribution to classification in the models in supplementary analysis (Supplementary Methods, Results, and Supplementary Figure 4).

We found that as measured by AUROC and AP, discriminatory performance of the random forest, logistic regression with L1 and elastic net penalties, and gradient boosting models was robust, even when up to 95% of the training set was removed. These findings showed that a training set size as small as 70-350 samples can maintain high performance, depending on the model.

Our results from traditional model performance and robustness evaluations show that our best machine learning phenotyping algorithm used Logistic Regression with L1 penalty or AdaBoost classifiers trained with controls without any cerebrovascular disease-related codes and a stroke service case population. However, we found that a similar model performed comparably well using cases identified by AIS-related diagnosis codes, suggesting that these models do not require manual case curation for high performance. In addition, our validation study in the UK Biobank detected AIS patients without ICD10 codes up to 250-fold better than random selection. This study has several limitations. First, we relied on noisy labels and proxies for training our models, as evidenced by the false positive rate of 4-16% that was determined by manual review. Without a gold standard set of cases, model performance is difficult to definitively evaluate. Second, we used only structured features contained within standard terminologies across the patients’ entire timeline, and did not use clinical notes. While clinical notes may contain much highly relevant information, they may also give rise to less reproducible and generalizable feature sets. Additionally, each feature contributed incrementally to high performance of the models and required minimal processing to acquire. Third, due to limitations of time and computational complexity, we did not exhaustively explore all possible combinations of cases and controls, including other potential AIS mimetic diseases. Despite these limitations, precision in the internal validation using the held-out set was high, and when applied to an external validation cohort, the developed models improved detection of AIS patients between 65 and 250-fold over random patient identification. Fourth, we did not study clinical implementation of the models. However, the discriminatory ability of the classifiers in the external validation suggest that although these models have not been implemented clinically, they may potentially be useful for improving the power of existing clinical and research study cohorts.

Our study benefits from several strengths. First, to address the current deficiencies in developing phenotyping algorithms, we developed an approach that demonstrates comparable discriminatory ability of identifying patients with AIS to past methods but has the added benefit of using EHR data that is generally available during inpatient hospitalization. Second, our model features were composed of structured data that encompass a larger feature variety than purely ICD-code based algorithms. Third, because our model incorporated structured data from standard terminologies, they therefore may be generalizable to other health systems outside CUIMC, whereas recent studies have relied on manually curated feature sets.^15^ Fourth, we examined several different combinations of cases, controls and classifiers for the purposes of training phenotyping models. Fourth, our phenotype classifiers assign probability of having had an AIS, which moves beyond binary classification of patients to develop a more granular description of patient’s disease state.

## SUMMARY

In addition to research tasks such as cohort identification, future models could focus on timely interventions such as care planning prior to discharge and risk stratification. We showed that structured data may be sufficiently accurate for classification, allowing for widespread usability of the algorithm. We also demonstrated the potential for using machine learning classifiers for cohort identification, which achieve high performance with many features acquired through minimal processing. In addition, patient cohorts derived using AIS diagnosis codes may obviate the need for manually-curated cohorts of patients with AIS, and procedure codes may be useful in identifying patients with AIS that may not have been coded with AIS-related diagnosis codes. We, and others, hypothesize that expanding cohort size by assigning a probability of disease may improve the power of heritability and genome-wide association studies.^25–30^ Utilizing the structured framework present in many current EHRs, along with machine learning models may provide a generalizable approach for expanding research study cohort size.

## ACKNOWLEDGEMENTS

PMT and NPT designed the study and drafted the original manuscript; BRK and MSE provided list of stroke service patients; BRK and PMT performed chart evaluation of stroke service patients; PMT and NPT performed analyses and wrote code. TL wrote code; MSE and NPT provided supervision; and PMT, BRK, TL, MSE, and NPT provided critical feedback on the manuscript. This research has been conducted using the UK Biobank Resource under Application Number 41039. We thank Dr. Patrick Ryan, Dr. Theresa Koleck, Dr. Prashanth Selvaraj, Dr. Rami Vanguri, Dr. Joseph Romano, Alexandre Yahi, and Dr. Kayla Quinnies for their feedback and guidance.

