## SupplementalMaterials for "Comparative analysis, applications, and interpretation of electronic health record-based stroke phenotyping methods"

### SUPPLEMENTARY MATERIAL

#### SUPPLEMENTARY METHODS

##### Hyperparameters for models

We ran a grid search for the hyperparameters of maximum tree depth, number of estimators, L1-L2 ratio, penalty parameters, subsampling rate, learning rate, momentum, and dropout. Outside of the default parameters, we used a max tree depth of 100, 1000 estimators, and square root maximum feature number for the random forest models. For L1 Logistic regression, we used an inverse of regularization strength of 0.1, and for elastic net regularization we used a penalty parameter of 0.01 and L1-L2 ratio parameter of .01, and "log" loss. For the boosting algorithms, we used a learning rate of 0.1, 1000 estimators, and for gradient boosting we also subsampled at a rate of 0.5, max depth as 10, and square root maximum feature number. The neural network model was comprised of two layers, the first with 64 neurons, relu activator, and l1 kernel regularizer. The second layer contained two neurons and a softmax activator. Learning was compiled by stochastic gradient descent with a learning rate of 0.01 and momentum of 0.9, nesterov=True. We included a dropout of data in the first layer at a rate of 0.3. Loss was calculated by categorical cross-entropy.

##### robustness of model

To test the robustness of the models, we trained on 100%, 90%, 70%, 50%, 30%, 10%, 5%, 1%, 0.5%, and 0.1% of the original training set. We then evaluated the performance of each training size 10 times on the test set through AUROC and AP.

##### Calibration

The classifiers assign a probability of being an AIS case to each patient. In order to interpret the meaning of these model probabilities and their applicability to other patients, we developed an empirical calibration method for the models. For each training fold during 10-fold cross validation, we first sorted the probabilities by value. We then averaged a bin of 100 patients intervals based on their assigned probability (predicted probability). Within each bin, we calculated the proportion of cases (actual probability). We repeated this sliding bin averaging for every 10 patients, creating an empirical function between the training set predicted probabilities and actual probabilities averaged across training folds. We then repeated the same averaging of predicted and actual probabilities for the sorted test set. We calibrated the test set predicted probabilities using the empirical function generated from the training folds. To evaluate calibration success, particularly for identifying cases, we calculated the root mean squared error (RMSE) of all calibrated test probabilities larger than 0.1. We made this cut off because many of the poorly calibrated models have many points near zero, giving a false low RMSE score. Models without any calibrated probabilities larger than 0.1 were given an RMSE of 'N/A'.

##### Feature Collapsing

Reduction of feature size is also essential for reproducibility of the models in other hospital settings. Our approach collapses features together by utilizing the hierarchical structure of ICD9, ICD10, and CPT4 diagnosis codes and procedure codes, and ATC ingredient definitions for medications. We mapped ICD9 and ICD10 diagnosis and procedure codes and drug prescription order ingredients to

SNOMED CT concept IDs. About 40% of the features could not be mapped automatically to SNOMED concepts, so we kept codes and ingredients that did not have concept ids as separate features. We then mapped ICD9 codes to ICD10 codes using General Equivalence Mappings.[1] We mapped all ICD9 and ICD10 diagnosis and procedure codes and CPT4 procedure codes to the Clinical Classifications System defined by the Health Cost and Utilization Project.[2] The CCS has flat level codes (most granular), and multilevel codes level 1-3 (least to most granular). We mapped all codes to the flat, multilevel level 1, and multilevel level 2 codes. We then mapped drug prescription ingredients to ATC codes by matching corresponding RxNorm codes to ATC codes. Some drugs required manual mapping to ATC codes. All in all, we successfully mapped 70% of 45 million prescription orders to ATC codes. We then reduced prescription order features to the five different levels of ATC description, which include, in increasing order of specificity, anatomical systems (14), therapeutic subgroups (78), pharmacological subgroups (159), chemical subgroups (364), and chemical substances (895). We then reran the training sets with collapsed features in three combinations: flat CCS level with chemical substrates (the most granular), multilevel CCS level 2 with pharmacological subgroups, and multilevel CCS level 1 with anatomical systems (the least granular). We chose these pairings to maintain similar ratios of drugs and ICD9 and ICD10 codes. We evaluated the robustness of hierarchical feature collapsing using the method described above. We then compared performance across levels of granularity using AUROC and average precision score, which is a proxy for area under the precision-recall curve.

**generalized linear model to determine feature category contribution to models**

To further explore the contribution of conditions, procedures, drugs, and demographics, to the models' classification performance, we performed a multivariate generalized linear model (GLM) on the stroke probabilities generated from each feature category. We retrained the models solely using features from each category. We also added a fifth category, which comprised the ICD9 codes that make up the T-L criteria and ICD10 equivalent codes (see Table 1). We added this category to compare the performance of the T-L criteria to the other features in AIS patient classification because our models do not include the T-L criteria in training. We then generated AIS probabilities for the holdout test set using each of the retrained models. Finally, we combined the model probabilities from each of the five models and ran the generalized linear model with binomial distribution to determine which categories significantly ($p<.05$) contributed to case determination and their corresponding beta coefficients.

#### SUPPLEMENTARY TABLES

| Abbreviation | Meaning |
|---|---|
| **AIS** | Acute Ischemic Stroke |
| **T-L** | Tirschwell- Longstreth |
| **ICD9** | International Classification of Disease, Version 9 |
| **ICD10** | International Classification of Disease, Version 10 |
| **ATC** | Anatomical Therapeutic Chemical Classification System |
| **CCS** | Clinical Classifications Software |
| **LR** | Logistic Regression with L1 penalty |
| **EN** | Logistic Regression with Elastic Net penalty |
| **RF** | Random Forest |
| **AB** | Adaboost |
| **GB** | Gradient Boosting |
| **NN** | Neural Network |
| **CvD** | Cerebrovascular Disease |
| **SN** | Stroke Service Cases, Stroke Mimetic Controls |
| **SI** | Stroke Service Cases, Controls without T-L codes for AIS |
| **SC** | Stroke Service Cases, Controls without ICD9 or ICD10 codes for CvD |
| **SCI** | Stroke Service Cases, Controls with ICD9 or ICD10 codes for CvD, and without T-L codes for AIS |
| **SR** | Stroke Service Cases, Random patients in the EHR as controls |
| **TN** | Cases with T-L codes for AIS, Stroke Mimetic Controls |
| **TI** | Cases with T-L codes for AIS, Controls without T-L codes for AIS |
| **TC** | Cases with T-L codes for AIS, Controls without ICD9 or ICD10 codes for CvD |
| **TCI** | Cases with T-L codes for AIS, Controls with ICD9 or ICD10 codes for CvD, and without T-L codes for AIS |
| **TR** | Cases with T-L codes for AIS, Random patients in the EHR as controls |
| **CN** | Cases with ICD9 or ICD10 codes for CvD, Stroke Mimetic Controls |
| **CI** | Cases with ICD9 or ICD10 codes for CvD, Controls without T-L codes for AIS |
| **CC** | Cases with ICD9 or ICD10 codes for CvD, Controls without ICD9 or ICD10 codes for CvD |
| **CCI** | Cases with ICD9 or ICD10 codes for CvD, Controls with ICD9 or ICD10 codes for CvD and without T-L codes for AIS |
| **CR** | Cases with ICD9 or ICD10 codes for CvD, Random patients in the EHR as controls |
| **RMSE** | Root Mean Squared Error |

| | |
|---|---|
| **GLM** | Generalized Linear Model |
| **CMS** | Centers for Medicare and Medicaid Services |
| **AUROC** | Area under the Receiver Operating Curve |
| **AP** | Average Precision Score |
| **P** | Precision |
| **R** | Recall |
| **FB** | F-score with beta coefficient B |
| **EHR** | Electronic Health Record |
| **PPV** | Positive Predictive Value |
| **NPV** | Negative Predictive Value |
| **CUIMC** | Columbia University Irving Medical Center |
| **CDW** | Common Data Warehouse |

**Supplementary Table 1: Abbreviations for Study**

| Variable | Case- S | Case- T | Case- C |
|---|---|---|---|
| N | 3473 | 6376-8000 | 6271-8000 |
| Gender, Female | 1709 (49.2%) | 3351-4222 (51.8-52.8%) | 3298-4278 (51.9-53.1%) |
| Age > 50 | 3115 (89.7%) | 5367-6747 (83.6-84.3%) | 5194-6570 (81.4-82.8%) |
| Race/Ethnicity | | | |
| Black or African American | 283 (8.15%) | 330-348 (4.18-5.18%) | 268-355 (3.94-4.43%) |
| White | 742 (21.4%) | 1083-1364 (16.2-17.0%) | 1263-1654 (19.7-20.7%) |
| Hispanic or Latino | 597 (17.2%) | 496-677 (7.78-8.46%) | 469-653 (7.48-8.16%) |
| Other | 40 (1.15%) | 43-69 (0.588%-0.862%) | 64-76 (0.825-1.02%) |
| Unknown/Declined to answer | 1811(52.1%) | 4425-5646 (69.4-70.6%) | 4204-5431 (66.6-67.9%) |

| | Control- N | Control- C | Control- I | Control- CI | Control- R |
|---|---|---|---|---|---|
| N | 3465-6346 | 3473-8000 | 3473-8000 | 3473-8000 | 3458-7920 |
| Gender, Female | 2233-4081 (64.3-64.4%) | 1927-4492 (55.5-56.1%) | 1945-4493 (55.4-56.2%) | 1828-4298 (52.6-53.7%) | 1970-4547 (56.2-57.4%) |
| Age > 50 | 1746-3225 (50.4-50.8%) | 1029-2366 (29.4-29.6%) | 1040-2458 (29.9-30.7%) | 2781-6454 (80.0-80.7%) | 1950-4524 (56.4-57.1%) |
| Race/Ethnicity | | | | | |
| Black or African American | 145-277 (4.18-4.36%) | 116-271 (3.26-3.39%) | 129-295 (3.54-3.71%) | 130-342 (3.74-4.28%) | 200-495 (5.78-6.27%) |
| White | 591-1113 (17.1-17.5%) | 300-734 (8.50-9.18%) | 348-725 (8.76-10.0%) | 759-1848 (21.8-23.0%) | 814-2021 (23.5-25.5%) |
| Hispanic or Latino | 276-482 (7.60-7.96%) | 232-559 (6.51-6.99%) | 233-515 (5.79-6.70%) | 283-630 (7.10-8.15%) | 436-1111 (12.6-14.0%) |
| Other | 30-38 (0.599-0.866%) | 28-60 (0.662-0.806%) | 29-71 (0.738-0.888%) | 33-87 (0.950-1.09%) | 65-144 (1.75-1.88%) |

| | | | | | |
|---|---|---|---|---|---|
| **Unknown/Declined to answer** | 2424-4440 (70.0%) | 2789-6470 (79.7-80.9%) | 2734-6470 (78.7-80.9%) | 2265-5155 (64.4-65.2%) | 1943-4293 (52.8-56.2%) |

**Supplementary Table 2:** Demographics of Case-control cohorts.

| Case/Control Combo | LR AUROC | RF AUROC | AB AUROC | GB AUROC | EN AUROC | LR Ave. Precision | RF Ave. Precision | AB Ave. Precision | GB Ave. Precision | EN Ave. Precision |
|---|---|---|---|---|---|---|---|---|---|---|
| **SN** | 0.995 | 0.995 | 0.994 | 0.951 | 0.996 | 0.939 | 0.843 | 0.936 | 0.760 | 0.933 |
| **SI** | 0.996 | 0.996 | 0.996 | 0.995 | 0.996 | 0.946 | 0.889 | 0.956 | 0.959 | 0.941 |
| **SC** | 0.997 | 0.996 | 0.996 | 0.977 | 0.996 | 0.943 | 0.894 | 0.963 | 0.818 | 0.936 |
| **SCI** | 0.996 | 0.996 | 0.995 | 0.963 | 0.997 | 0.969 | 0.956 | 0.964 | 0.796 | 0.964 |
| **SR** | 0.995 | 0.995 | 0.993 | 0.986 | 0.996 | 0.966 | 0.919 | 0.961 | 0.936 | 0.969 |
| **TN** | 0.967 | 0.982 | 0.969 | 0.956 | 0.962 | 0.814 | 0.461 | 0.755 | 0.641 | 0.770 |
| **TI** | 0.995 | 0.996 | 0.993 | 0.994 | 0.997 | 0.930 | 0.917 | 0.845 | 0.844 | 0.945 |
| **TC** | 0.994 | 0.996 | 0.995 | 0.995 | 0.998 | 0.935 | 0.907 | 0.932 | 0.867 | 0.948 |
| **TCI** | 0.943 | 0.979 | 0.930 | 0.942 | 0.953 | 0.836 | 0.865 | 0.577 | 0.673 | 0.861 |
| **TR** | 0.923 | 0.837 | 0.865 | 0.828 | 0.900 | 0.720 | 0.332 | 0.500 | 0.542 | 0.717 |
| **CN** | 0.958 | 0.965 | 0.947 | 0.890 | 0.949 | 0.729 | 0.183 | 0.513 | 0.431 | 0.674 |
| **CI** | 0.995 | 0.995 | 0.995 | 0.993 | 0.997 | 0.916 | 0.811 | 0.874 | 0.814 | 0.930 |
| **CC** | 0.995 | 0.996 | 0.995 | 0.994 | 0.997 | 0.918 | 0.866 | 0.895 | 0.843 | 0.931 |
| **CCI** | 0.950 | 0.961 | 0.902 | 0.866 | 0.954 | 0.837 | 0.851 | 0.336 | 0.477 | 0.838 |
| **CR** | 0.913 | 0.771 | 0.858 | 0.784 | 0.907 | 0.662 | 0.258 | 0.369 | 0.448 | 0.679 |

**Supplementary Table 3:** Area under the receiver-operating characteristic curve and average precision score across models for holdout test set. (LR) logistic regression with l1 penalty, (RF) random forest, (AB) AdaBoost, (GB) gradient boosting, (EN) logistic regression with elastic net penalty. Different combinations of cases and controls are shown on the y-axis. Cases (first letter) may be one of cerebrovascular (C), T-L (T), or Stroke Service (S), see Methods "Data" for definition of case sets. Controls (second and third letters) may be one of random (R), cerebrovascular disease but no AIS code (CI), no cerebrovascular disease (C), no AIS code (I), or a stroke mimetic disease (N), see Method "Data"

| Case/ Control | LR Sens | RF Sens | AB Sens | GB Sens | EN Sens | LR Spec | RF Spec | AB Spec | GB Spec | EN Spec | LR PPV | RF PPV | AB PPV | GB PPV | EN PPV | LR NPV | RF NPV | AB NPV | GB NPV | EN NPV |
|---|---|---|---|---|---|---|---|---|---|---|---|---|---|---|---|---|---|---|---|---|
| **SN** | 0.749 | 0.522 | 0.753 | 0.699 | 0.715 | 1.000 | 0.999 | 1.000 | 0.996 | 0.999 | 0.007 | 0.005 | 0.007 | 0.007 | 0.007 | 0.997 | 0.995 | 0.998 | 0.997 | 0.997 |
| **SI** | 0.842 | 0.748 | 0.923 | 0.766 | 0.825 | 0.998 | 0.998 | 0.997 | 0.996 | 0.998 | 0.008 | 0.007 | 0.009 | 0.008 | 0.008 | 0.998 | 0.997 | 0.999 | 0.998 | 0.998 |
| **SC** | 0.941 | 0.791 | 0.959 | 0.836 | 0.902 | 0.996 | 0.997 | 0.994 | 0.996 | 0.996 | 0.009 | 0.008 | 0.010 | 0.008 | 0.009 | 0.999 | 0.998 | 1.000 | 0.998 | 0.999 |
| **SCI** | 0.575 | 0.450 | 0.524 | 0.428 | 0.651 | 1.000 | 1.000 | 1.000 | 0.998 | 1.000 | 0.006 | 0.004 | 0.005 | 0.004 | 0.006 | 0.996 | 0.995 | 0.995 | 0.994 | 0.997 |
| **SR** | 0.586 | 0.549 | 0.546 | 0.522 | 0.643 | 1.000 | 1.000 | 1.000 | 1.000 | 1.000 | 0.006 | 0.005 | 0.005 | 0.005 | 0.006 | 0.996 | 0.996 | 0.995 | 0.995 | 0.996 |
| **TN** | 0.525 | 0.331 | 0.461 | 0.434 | 0.388 | 1.000 | 0.997 | 0.999 | 0.998 | 1.000 | 0.005 | 0.003 | 0.005 | 0.004 | 0.004 | 0.995 | 0.993 | 0.995 | 0.994 | 0.994 |
| **TI** | 0.743 | 0.692 | 0.576 | 0.614 | 0.705 | 0.999 | 0.999 | 0.999 | 0.999 | 1.000 | 0.007 | 0.007 | 0.006 | 0.006 | 0.007 | 0.997 | 0.997 | 0.996 | 0.996 | 0.997 |
| **TC** | 0.822 | 0.737 | 0.827 | 0.784 | 0.781 | 0.999 | 0.999 | 0.999 | 0.998 | 0.999 | 0.008 | 0.007 | 0.008 | 0.008 | 0.008 | 0.998 | 0.997 | 0.998 | 0.998 | 0.998 |
| **TCI** | 0.205 | 0.381 | 0.329 | 0.439 | 0.250 | 1.000 | 1.000 | 0.997 | 0.999 | 1.000 | 0.002 | 0.004 | 0.003 | 0.004 | 0.002 | 0.992 | 0.994 | 0.993 | 0.994 | 0.993 |
| **TR** | 0.602 | 0.248 | 0.522 | 0.370 | 0.552 | 0.997 | 0.999 | 0.996 | 0.998 | 0.999 | 0.006 | 0.002 | 0.005 | 0.004 | 0.005 | 0.996 | 0.993 | 0.995 | 0.994 | 0.996 |
| **CN** | 0.471 | 0.181 | 0.394 | 0.273 | 0.382 | 0.999 | 0.993 | 0.998 | 0.999 | 0.999 | 0.005 | 0.002 | 0.004 | 0.003 | 0.004 | 0.995 | 0.992 | 0.994 | 0.993 | 0.994 |
| **CI** | 0.736 | 0.746 | 0.706 | 0.667 | 0.752 | 0.999 | 0.996 | 0.998 | 0.998 | 0.999 | 0.007 | 0.007 | 0.007 | 0.007 | 0.007 | 0.997 | 0.997 | 0.997 | 0.997 | 0.998 |
| **CC** | 0.833 | 0.827 | 0.797 | 0.780 | 0.798 | 0.998 | 0.995 | 0.998 | 0.998 | 0.999 | 0.008 | 0.008 | 0.008 | 0.008 | 0.008 | 0.998 | 0.998 | 0.998 | 0.998 | 0.998 |
| **CCI** | 0.347 | 0.486 | 0.496 | 0.352 | 0.395 | 1.000 | 1.000 | 0.992 | 0.998 | 1.000 | 0.003 | 0.005 | 0.005 | 0.004 | 0.004 | 0.994 | 0.995 | 0.995 | 0.994 | 0.994 |
| **CR** | 0.596 | 0.274 | 0.538 | 0.424 | 0.649 | 0.995 | 0.996 | 0.995 | 0.997 | 0.982 | 0.006 | 0.003 | 0.005 | 0.004 | 0.007 | 0.996 | 0.993 | 0.995 | 0.994 | 0.996 |

**Supplementary Table 4:** Sensitivity, specificity, positive predictive value, and negative predictive value of models on holdout test set. See Supplementary Table 1 for case-control, model, and evaluator abbreviations' definitions.

| Case/ Control Combo | LR P at 50 | RF P at 50 | AB P at 50 | GB P at 50 | EN P at 50 | LR P at 100 | RF P at 100 | AB P at 100 | GB P at 100 | EN P at 100 | LR P at 500 | RF P at 500 | AB P at 500 | GB P at 500 | EN P at 500 | LR P at N Cases | RF P at N Cases | AB P at N Cases | GB P at N Cases | EN P at N Cases |
|---|---|---|---|---|---|---|---|---|---|---|---|---|---|---|---|---|---|---|---|---|
| **SN** | 1.00 | 1.00 | 1.00 | 1.00 | 0.86 | 1.00 | 1.00 | 1.00 | 0.92 | 0.75 | 1.00 | 1.00 | 1.00 | 0.99 | 0.87 | 0.76 | 0.79 | 0.79 | 0.79 | 0.67 |
| **SI** | 1.00 | 1.00 | 1.00 | 1.00 | 0.89 | 1.00 | 1.00 | 1.00 | 0.97 | 0.81 | 1.00 | 1.00 | 1.00 | 0.99 | 0.90 | 0.97 | 0.99 | 1.00 | 1.00 | 0.91 |
| **SC** | 1.00 | 1.00 | 1.00 | 0.99 | 0.88 | 1.00 | 1.00 | 1.00 | 0.97 | 0.81 | 1.00 | 1.00 | 1.00 | 1.00 | 0.91 | 0.80 | 0.80 | 0.80 | 0.80 | 0.73 |
| **SCI** | 1.00 | 1.00 | 1.00 | 1.00 | 0.92 | 1.00 | 1.00 | 1.00 | 1.00 | 0.90 | 1.00 | 1.00 | 1.00 | 1.00 | 0.91 | 0.72 | 0.77 | 0.79 | 0.81 | 0.78 |
| **SR** | 1.00 | 1.00 | 1.00 | 1.00 | 0.91 | 1.00 | 1.00 | 1.00 | 0.99 | 0.83 | 1.00 | 1.00 | 1.00 | 1.00 | 0.91 | 1.00 | 1.00 | 1.00 | 1.00 | 0.89 |
| **TN** | 1.00 | 1.00 | 1.00 | 0.98 | 0.76 | 0.68 | 0.72 | 0.70 | 0.58 | 0.49 | 1.00 | 1.00 | 0.98 | 0.89 | 0.74 | 0.90 | 0.95 | 0.94 | 0.81 | 0.63 |
| **TI** | 1.00 | 1.00 | 1.00 | 0.99 | 0.87 | 1.00 | 1.00 | 1.00 | 0.97 | 0.84 | 0.99 | 1.00 | 0.99 | 0.92 | 0.77 | 1.00 | 0.99 | 0.98 | 0.92 | 0.77 |
| **TC** | 1.00 | 1.00 | 1.00 | 0.99 | 0.88 | 1.00 | 1.00 | 1.00 | 0.96 | 0.83 | 1.00 | 1.00 | 1.00 | 0.99 | 0.88 | 1.00 | 0.99 | 0.98 | 0.92 | 0.81 |
| **TCI** | 1.00 | 1.00 | 1.00 | 1.00 | 0.79 | 1.00 | 0.99 | 0.99 | 0.97 | 0.80 | 0.74 | 0.64 | 0.79 | 0.62 | 0.64 | 0.98 | 0.97 | 0.96 | 0.84 | 0.65 |
| **TR** | 1.00 | 1.00 | 1.00 | 0.96 | 0.68 | 0.99 | 0.97 | 0.88 | 0.58 | 0.39 | 0.83 | 0.73 | 0.73 | 0.73 | 0.55 | 0.99 | 0.99 | 0.98 | 0.82 | 0.54 |
| **CN** | 1.00 | 1.00 | 1.00 | 0.90 | 0.69 | 0.13 | 0.15 | 0.15 | 0.18 | 0.21 | 0.9 | 0.81 | 0.72 | 0.66 | 0.57 | 0.95 | 0.90 | 0.86 | 0.64 | 0.47 |
| **CI** | 1.00 | 1.00 | 1.00 | 0.99 | 0.85 | 1.00 | 1.00 | 0.99 | 0.89 | 0.73 | 1.00 | 1.00 | 0.99 | 0.95 | 0.80 | 0.97 | 0.98 | 0.97 | 0.89 | 0.76 |
| **CC** | 1.00 | 1.00 | 1.00 | 0.99 | 0.84 | 1.00 | 1.00 | 1.00 | 0.93 | 0.79 | 1.00 | 1.00 | 1.00 | 0.98 | 0.82 | 0.95 | 0.96 | 0.97 | 0.93 | 0.78 |
| **CCI** | 1.00 | 1.00 | 1.00 | 0.98 | 0.78 | 1.00 | 1.00 | 1.00 | 0.98 | 0.81 | 0.63 | 0.70 | 0.69 | 0.21 | 0.32 | 0.99 | 0.96 | 0.91 | 0.69 | 0.50 |
| **CR** | 1.00 | 1.00 | 1.00 | 0.88 | 0.64 | 1.00 | 0.88 | 0.82 | 0.48 | 0.35 | 0.71 | 0.43 | 0.35 | 0.58 | 0.53 | 0.99 | 0.98 | 0.95 | 0.71 | 0.49 |

**Supplementary Table 5:** Precision at top 50, 100, and number of known cases for each classifier.

| | | Logistic Regression (L1 Penalty) | | | | Random Forest | | | | Adaboost | | | | Gradient Boosting | | | | Logistic Regression (Elastic Net Penalty) | | | |
|---|---|---|---|---|---|---|---|---|---|---|---|---|---|---|---|---|---|---|---|---|---|
| | **Metric** | **FB=1** | **FB=1/2** | **FB=1/4** | **FB=1/8** | **B=1** | **B=1/2** | **B=1/4** | **B=1/8** | **B=1** | **B=1/2** | **B=1/4** | **B=1/8** | **B=1** | **B=1/2** | **B=1/4** | **B=1/8** | **B=1** | **B=1/2** | **B=1/4** | **B=1/8** |
| **SN** | **P** | 0.97 (0.03) | 0.98 (0.03) | 0.98 (0.03) | 0.98 (0.03) | 0.90 (0.06) | 0.93 (0.06) | 0.93 (0.06) | 0.93 (0.06) | 0.90 (0.06) | 0.93 (0.06) | 0.93 (0.06) | 0.93 (0.06) | 0.95 (0.04) | 0.96 (0.05) | 0.96 (0.05) | 0.96 (0.05) | 0.85 (0.23) | 0.88 (0.24) | 0.88 (0.24) | 0.88 (0.24) |
| | **R** | 0.75 (0.08) | 0.66 (0.16) | 0.66 (0.16) | 0.66 (0.16) | 0.52 (0.11) | 0.39 (0.19) | 0.39 (0.19) | 0.39 (0.19) | 0.52 (0.11) | 0.39 (0.19) | 0.39 (0.19) | 0.39 (0.19) | 0.75 (0.08) | 0.59 (0.25) | 0.59 (0.25) | 0.59 (0.25) | 0.70 (0.13) | 0.58 (0.23) | 0.58 (0.23) | 0.58 (0.23) |
| | **F** | 0.84 (0.04) | 0.77 (0.13) | 0.77 (0.13) | 0.77 (0.13) | 0.65 (0) | 0.52 (0.20) | 0.52 (0.20) | 0.52 (0.20) | 0.65 (0) | 0.52 (0.20) | 0.52 (0.20) | 0.52 (0.20) | 0.84 (0.04) | 0.69 (0.22) | 0.69 (0.22) | 0.69 (0.22) | 0.74 (0.17) | 0.63 (0.26) | 0.63 (0.26) | 0.63 (0.26) |
| | **T** | 0.96 (0.03) | 0.97 (0.03) | 0.97 (0.03) | 0.97 (0.03) | 0.88 (0) | 0.90 (0.03) | 0.90 (0.03) | 0.90 (0.03) | 0.88 (0) | 0.90 (0.03) | 0.90 (0.03) | 0.90 (0.03) | 0.51 (0.00) | 0.52 (0.01) | 0.52 (0.01) | 0.52 (0.01) | 1.00 (0.00) | 1.00 (0.00) | 1.00 (0.00) | 1.00 (0.00) |
| | **# >T** | 773 (97) | 675 (176) | 675 (176) | 675 (176) | 591 (166) | 431 (226) | 431 (226) | 431 (226) | 795 (115) | 623 (285) | 623 (285) | 623 (285) | 1057 (852) | 904 (921) | 904 (921) | 904 (921) | 777 (201) | 702 (240) | 702 (240) | 702 (240) |
| | **# > T in EHR** | 37942 | | | | 37911 | | | | 54587 | | | | 67459 | | | | 84222 | | | |
| **SI** | **P** | 0.87 (0.11) | 0.87 (0.11) | 0.87 (0.11) | 0.87 (0.11) | 0.84 (0.08) | 0.92 (0.09) | 0.92 (0.09) | 0.92 (0.09) | 0.84 (0.08) | 0.92 (0.09) | 0.92 (0.09) | 0.92 (0.09) | 0.78 (0.13) | 0.82 (0.14) | 0.82 (0.14) | 0.82 (0.14) | 0.78 (0.27) | 0.79 (0.28) | 0.79 (0.28) | 0.79 (0.28) |
| | **R** | 0.84 (0.08) | 0.81 (0.12) | 0.81 (0.12) | 0.81 (0.12) | 0.75 (0.12) | 0.53 (0.25) | 0.53 (0.25) | 0.53 (0.25) | 0.75 (0.12) | 0.53 (0.25) | 0.53 (0.25) | 0.53 (0.25) | 0.92 (0.05) | 0.77 (0.32) | 0.77 (0.32) | 0.77 (0.32) | 0.77 (0.17) | 0.70 (0.25) | 0.70 (0.25) | 0.70 (0.25) |
| | **F** | 0.84 (0.03) | 0.83 (0.05) | 0.83 (0.05) | 0.83 (0.05) | 0.78 (0) | 0.62 (0.20) | 0.62 (0.20) | 0.62 (0.20) | 0.78 (0) | 0.62 (0.20) | 0.62 (0.20) | 0.62 (0.20) | 0.84 (0.05) | 0.72 (0.24) | 0.72 (0.24) | 0.72 (0.24) | 0.75 (0.21) | 0.69 (0.25) | 0.69 (0.25) | 0.69 (0.25) |
| | **T** | 0.93 (0.08) | 0.93 (0.08) | 0.93 (0.08) | 0.93 (0.08) | 0.93 (0) | 0.96 (0.02) | 0.96 (0.02) | 0.96 (0.02) | 0.93 (0) | 0.96 (0.02) | 0.96 (0.02) | 0.96 (0.02) | 0.51 (0.01) | 0.52 (0.02) | 0.52 (0.02) | 0.52 (0.02) | 1.00 (0.00) | 1.00 (0.00) | 1.00 (0.00) | 1.00 (0.00) |
| | **# >T** | 999 (233) | 964 (269) | 964 (269) | 964 (269) | 909 (210) | 612 (336) | 612 (336) | 612 (336) | 1220 (256) | 1022 (495) | 1022 (495) | 1022 (495) | 1168 (700) | 1100 (760) | 1100 (760) | 1100 (760) | 980 (228) | 910 (212) | 910 (212) | 910 (212) |
| | **# > T in EHR** | 57057 | | | | 104710 | | | | 79922 | | | | 92435 | | | | 110097 | | | |
| **SC** | **P** | 0.70 (0.11) | 0.70 (0.11) | 0.70 (0.11) | 0.70 (0.11) | 0.76 (0.14) | 0.79 (0.16) | 0.79 (0.16) | 0.79 (0.16) | 0.76 (0.14) | 0.79 (0.16) | 0.79 (0.16) | 0.79 (0.16) | 0.67 (0.16) | 0.68 (0.17) | 0.68 (0.17) | 0.68 (0.17) | 0.77 (0.23) | 0.77 (0.23) | 0.77 (0.23) | 0.77 (0.23) |
| | **R** | 0.94 (0.03) | 0.94 (0.03) | 0.94 (0.03) | 0.94 (0.03) | 0.79 (0.15) | 0.71 (0.24) | 0.71 (0.24) | 0.71 (0.24) | 0.79 (0.15) | 0.71 (0.24) | 0.71 (0.24) | 0.71 (0.24) | 0.96 (0.03) | 0.93 (0.11) | 0.93 (0.11) | 0.93 (0.11) | 0.84 (0.16) | 0.84 (0.16) | 0.84 (0.16) | 0.84 (0.16) |

| | | | | | | | | | | | | | | | | | | | | | |
|---|---|---|---|---|---|---|---|---|---|---|---|---|---|---|---|---|---|---|---|---|---|
| | **F** | 0.79 (0.06) | 0.79 (0.06) | 0.79 (0.06) | 0.79 (0.06) | 0.75 (0) | 0.69 (0.12) | 0.69 (0.12) | 0.69 (0.12) | 0.75 (0) | 0.69 (0.12) | 0.69 (0.12) | 0.69 (0.12) | 0.77 (0.10) | 0.76 (0.09) | 0.76 (0.09) | 0.76 (0.09) | 0.78 (0.16) | 0.78 (0.16) | 0.78 (0.16) | 0.78 (0.16) |
| | **T** | 0.82 (0.11) | 0.82 (0.11) | 0.82 (0.11) | 0.82 (0.11) | 0.90 (0) | 0.91 (0.05) | 0.91 (0.05) | 0.91 (0.05) | 0.90 (0) | 0.91 (0.05) | 0.91 (0.05) | 0.91 (0.05) | 0.51 (0.01) | 0.51 (0.02) | 0.51 (0.02) | 0.51 (0.02) | 0.98 (0.05) | 0.98 (0.05) | 0.98 (0.05) | 0.98 (0.05) |
| | **# >T** | 1386 (253) | 1386 (253) | 1386 (253) | 1386 (253) | 1108 (385) | 1003 (492) | 1003 (492) | 1003 (492) | 1523 (399) | 1486 (459) | 1486 (459) | 1486 (459) | 1197 (414) | 1197 (414) | 1197 (414) | 1197 (414) | 1332 (393) | 1332 (393) | 1332 (393) | 1332 (393) |
| | **# > T in EHR** | 67350 | | | | 91099 | | | | 81257 | | | | 94066 | | | | 109483 | | | |
| | **P** | 1.00 (0.00) | 1.00 (0.00) | 1.00 (0.00) | 1.00 (0.00) | 1.00 (0.00) | 1.00 (0.00) | 1.00 (0.00) | 1.00 (0.00) | 1.00 (0.00) | 1.00 (0.00) | 1.00 (0.00) | 1.00 (0.00) | 1.00 (0.00) | 1.00 (0.00) | 1.00 (0.00) | 1.00 (0.00) | 0.91 (0.23) | 0.91 (0.24) | 0.91 (0.24) | 0.91 (0.24) |
| | **R** | 0.57 (0.10) | 0.45 (0.14) | 0.45 (0.14) | 0.45 (0.14) | 0.45 (0.14) | 0.28 (0.10) | 0.28 (0.10) | 0.28 (0.10) | 0.45 (0.14) | 0.28 (0.10) | 0.28 (0.10) | 0.28 (0.10) | 0.52 (0.10) | 0.47 (0.10) | 0.47 (0.10) | 0.47 (0.10) | 0.43 (0.09) | 0.41 (0.08) | 0.41 (0.08) | 0.41 (0.08) |
| | **F** | 0.72 (0.08) | 0.61 (0.15) | 0.61 (0.15) | 0.61 (0.15) | 0.61 (0) | 0.43 (0.11) | 0.43 (0.11) | 0.43 (0.11) | 0.61 (0) | 0.43 (0.11) | 0.43 (0.11) | 0.43 (0.11) | 0.68 (0.09) | 0.64 (0.09) | 0.64 (0.09) | 0.64 (0.09) | 0.55 (0.12) | 0.54 (0.12) | 0.54 (0.12) | 0.54 (0.12) |
| **SCI** | **T** | 0.94 (0.03) | 0.97 (0.02) | 0.97 (0.02) | 0.97 (0.02) | 0.84 (0) | 0.89 (0.02) | 0.89 (0.02) | 0.89 (0.02) | 0.84 (0) | 0.89 (0.02) | 0.89 (0.02) | 0.89 (0.02) | 0.51 (0.00) | 0.51 (0.00) | 0.51 (0.00) | 0.51 (0.00) | 1.00 (0.00) | 1.00 (0.00) | 1.00 (0.00) | 1.00 (0.00) |
| | **# >T** | 575 (104) | 455 (143) | 455 (143) | 455 (143) | 450 (139) | 280 (94) | 280 (94) | 280 (94) | 526 (105) | 476 (103) | 476 (103) | 476 (103) | 622 (610) | 601 (599) | 601 (599) | 601 (599) | 654 (92) | 503 (93) | 503 (93) | 503 (93) |
| | **# > T in EHR** | 10052 | | | | 6464 | | | | 12647 | | | | 14302 | | | | 20033 | | | |
| | **P** | 0.99 (0.01) | 1.00 (0.00) | 1.00 (0.00) | 1.00 (0.00) | 0.98 (0.02) | 0.98 (0.02) | 0.98 (0.02) | 0.98 (0.02) | 0.98 (0.02) | 0.98 (0.02) | 0.98 (0.02) | 0.98 (0.02) | 1.00 (0.00) | 1.00 (0.00) | 1.00 (0.00) | 1.00 (0.00) | 1.00 (0.00) | 1.00 (0.00) | 1.00 (0.00) | 1.00 (0.00) |
| | **R** | 0.59 (0.15) | 0.29 (0.20) | 0.29 (0.20) | 0.29 (0.20) | 0.55 (0.12) | 0.45 (0.17) | 0.45 (0.17) | 0.45 (0.17) | 0.55 (0.12) | 0.45 (0.17) | 0.45 (0.17) | 0.45 (0.17) | 0.55 (0.14) | 0.34 (0.13) | 0.34 (0.13) | 0.34 (0.13) | 0.52 (0.17) | 0.35 (0.17) | 0.35 (0.17) | 0.35 (0.17) |
| | **F** | 0.73 (0.12) | 0.41 (0.23) | 0.41 (0.23) | 0.41 (0.23) | 0.69 (0) | 0.60 (0.16) | 0.60 (0.16) | 0.60 (0.16) | 0.69 (0) | 0.60 (0.16) | 0.60 (0.16) | 0.60 (0.16) | 0.69 (0.12) | 0.49 (0.14) | 0.49 (0.14) | 0.49 (0.14) | 0.67 (0.17) | 0.49 (0.19) | 0.49 (0.19) | 0.49 (0.19) |
| **SR** | **T** | 0.94 (0.05) | 0.99 (0.01) | 0.99 (0.01) | 0.99 (0.01) | 0.84 (0) | 0.87 (0.05) | 0.87 (0.05) | 0.87 (0.05) | 0.84 (0) | 0.87 (0.05) | 0.87 (0.05) | 0.87 (0.05) | 0.51 (0.00) | 0.52 (0.00) | 0.52 (0.00) | 0.52 (0.00) | 1.00 (0.00) | 1.00 (0.00) | 1.00 (0.00) | 1.00 (0.00) |
| | **# >T** | 589 (151) | 287 (196) | 287 (196) | 287 (196) | 563 (129) | 461 (185) | 461 (185) | 461 (185) | 548 (144) | 338 (133) | 338 (133) | 338 (133) | 524 (170) | 345 (171) | 345 (171) | 345 (171) | 648 (132) | 378 (223) | 378 (223) | 378 (223) |
| | **# > T in EHR** | 13261 | | | | 11569 | | | | 18079 | | | | 25379 | | | | 36925 | | | |

| | | | | | | | | | | | | | | | | | | | | | |
|---|---|---|---|---|---|---|---|---|---|---|---|---|---|---|---|---|---|---|---|---|---|
| | **P** | 0.97 (0.02) | 1.00 (0.00) | 1.00 (0.00) | 1.00 (0.00) | 0.56 (0.07) | 0.70 (0.14) | 0.70 (0.14) | 0.70 (0.14) | 0.56 (0.07) | 0.70 (0.14) | 0.70 (0.14) | 0.70 (0.14) | 0.89 (0.02) | 0.99 (0.01) | 0.99 (0.01) | 0.99 (0.01) | 0.76 (0.08) | 0.84 (0.28) | 0.84 (0.28) | 0.84 (0.28) |
| | **R** | 0.52 (0.08) | 0.10 (0.05) | 0.10 (0.05) | 0.10 (0.05) | 0.33 (0.12) | 0.04 (0.03) | 0.04 (0.03) | 0.04 (0.03) | 0.33 (0.12) | 0.04 (0.03) | 0.04 (0.03) | 0.04 (0.03) | 0.46 (0.12) | 0.09 (0.05) | 0.09 (0.05) | 0.09 (0.05) | 0.43 (0.11) | 0.07 (0.08) | 0.07 (0.08) | 0.07 (0.08) |
| | **F** | 0.68 (0.07) | 0.18 (0.09) | 0.18 (0.09) | 0.18 (0.09) | 0.40 (0) | 0.07 (0.05) | 0.07 (0.05) | 0.07 (0.05) | 0.40 (0) | 0.07 (0.05) | 0.07 (0.05) | 0.07 (0.05) | 0.60 (0.11) | 0.15 (0.09) | 0.15 (0.09) | 0.15 (0.09) | 0.54 (0.06) | 0.13 (0.13) | 0.13 (0.13) | 0.13 (0.13) |
| **TN** | **T** | 0.92 (0.02) | 0.99 (0.01) | 0.99 (0.01) | 0.99 (0.01) | 0.81 (0) | 0.88 (0.02) | 0.88 (0.02) | 0.88 (0.02) | 0.81 (0) | 0.88 (0.02) | 0.88 (0.02) | 0.88 (0.02) | 0.51 (0.00) | 0.51 (0.00) | 0.51 (0.00) | 0.51 (0.00) | 0.98 (0.02) | 1.00 (0.00) | 1.00 (0.00) | 1.00 (0.00) |
| | **# >T** | 542 (96) | 99 (53) | 99 (53) | 99 (53) | 623 (311) | 59 (45) | 59 (45) | 59 (45) | 522 (152) | 86 (54) | 86 (54) | 86 (54) | 590 (236) | 80 (88) | 80 (88) | 80 (88) | 417 (200) | 14 (12) | 14 (12) | 14 (12) |
| | **# > T in EHR** | 45830 | | | | 44533 | | | | 47446 | | | | 52497 | | | | 136289 | | | |
| | **P** | 0.92 (0.05) | 0.99 (0.01) | 0.99 (0.01) | 0.99 (0.01) | 0.91 (0.06) | 1.00 (0.01) | 1.00 (0.01) | 1.00 (0.01) | 0.91 (0.06) | 1.00 (0.01) | 1.00 (0.01) | 1.00 (0.01) | 0.88 (0.05) | 0.97 (0.04) | 0.97 (0.04) | 0.97 (0.04) | 0.87 (0.03) | 0.96 (0.03) | 0.96 (0.03) | 0.96 (0.03) |
| | **R** | 0.74 (0.13) | 0.31 (0.20) | 0.31 (0.20) | 0.31 (0.20) | 0.69 (0.12) | 0.12 (0.10) | 0.12 (0.10) | 0.12 (0.10) | 0.69 (0.12) | 0.12 (0.10) | 0.12 (0.10) | 0.12 (0.10) | 0.58 (0.12) | 0.23 (0.14) | 0.23 (0.14) | 0.23 (0.14) | 0.61 (0.10) | 0.14 (0.11) | 0.14 (0.11) | 0.14 (0.11) |
| | **F** | 0.81 (0.06) | 0.44 (0.23) | 0.44 (0.23) | 0.44 (0.23) | 0.77 (0) | 0.21 (0.15) | 0.21 (0.15) | 0.21 (0.15) | 0.77 (0) | 0.21 (0.15) | 0.21 (0.15) | 0.21 (0.15) | 0.68 (0.10) | 0.34 (0.17) | 0.34 (0.17) | 0.34 (0.17) | 0.71 (0.06) | 0.23 (0.16) | 0.23 (0.16) | 0.23 (0.16) |
| **TI** | **T** | 0.96 (0.03) | 1.00 (0.00) | 1.00 (0.00) | 1.00 (0.00) | 0.90 (0) | 0.96 (0.01) | 0.96 (0.01) | 0.96 (0.01) | 0.90 (0) | 0.96 (0.01) | 0.96 (0.01) | 0.96 (0.01) | 0.51 (0.00) | 0.52 (0.01) | 0.52 (0.01) | 0.52 (0.01) | 0.99 (0.00) | 1.00 (0.00) | 1.00 (0.00) | 1.00 (0.00) |
| | **# >T** | 814 (184) | 311 (199) | 311 (199) | 311 (199) | 774 (187) | 125 (98) | 125 (98) | 125 (98) | 663 (164) | 237 (160) | 237 (160) | 237 (160) | 710 (139) | 145 (115) | 145 (115) | 145 (115) | 737 (193) | 87 (109) | 87 (109) | 87 (109) |
| | **# > T in EHR** | 83554 | | | | 124453 | | | | 73771 | | | | 248110 | | | | 886089 | | | |
| | **P** | 0.91 (0.04) | 0.98 (0.02) | 0.98 (0.02) | 0.98 (0.02) | 0.88 (0.04) | 0.97 (0.03) | 0.97 (0.03) | 0.97 (0.03) | 0.88 (0.04) | 0.97 (0.03) | 0.97 (0.03) | 0.97 (0.03) | 0.92 (0.03) | 0.97 (0.04) | 0.97 (0.04) | 0.97 (0.04) | 0.80 (0.07) | 0.92 (0.04) | 0.92 (0.04) | 0.92 (0.04) |
| | **R** | 0.82 (0.07) | 0.53 (0.17) | 0.53 (0.17) | 0.53 (0.17) | 0.74 (0.10) | 0.36 (0.24) | 0.36 (0.24) | 0.36 (0.24) | 0.74 (0.10) | 0.36 (0.24) | 0.36 (0.24) | 0.36 (0.24) | 0.83 (0.05) | 0.54 (0.22) | 0.54 (0.22) | 0.54 (0.22) | 0.78 (0.09) | 0.43 (0.18) | 0.43 (0.18) | 0.43 (0.18) |
| **TC** | **F** | 0.86 (0.03) | 0.67 (0.15) | 0.67 (0.15) | 0.67 (0.15) | 0.80 (0) | 0.47 (0.27) | 0.47 (0.27) | 0.47 (0.27) | 0.80 (0) | 0.47 (0.27) | 0.47 (0.27) | 0.47 (0.27) | 0.87 (0.02) | 0.66 (0.19) | 0.66 (0.19) | 0.66 (0.19) | 0.79 (0.03) | 0.56 (0.15) | 0.56 (0.15) | 0.56 (0.15) |
| | **T** | 0.97 (0.02) | 1.00 (0.01) | 1.00 (0.01) | 1.00 (0.01) | 0.91 (0) | 0.95 (0.02) | 0.95 (0.02) | 0.95 (0.02) | 0.91 (0) | 0.95 (0.02) | 0.95 (0.02) | 0.95 (0.02) | 0.51 (0.00) | 0.52 (0.01) | 0.52 (0.01) | 0.52 (0.01) | 0.99 (0.01) | 1.00 (0.00) | 1.00 (0.00) | 1.00 (0.00) |
| | **# >T** | 906 (112) | 542 (186) | 542 (186) | 542 (186) | 841 (142) | 377 (255) | 377 (255) | 377 (255) | 901 (86) | 562 (254) | 562 (254) | 562 (254) | 992 (205) | 474 (219) | 474 (219) | 474 (219) | 840 (144) | 231 (232) | 231 (232) | 231 (232) |

| | | | | | | | | | | | | | | | | | | | | | |
|---|---|---|---|---|---|---|---|---|---|---|---|---|---|---|---|---|---|---|---|---|---|
| | **# > T in EHR** | 92041 | | | | 141147 | | | | 105523 | | | | 86548 | | | | 143860 | | | |
| | **P** | 1.00 (0.00) | 1.00 (0.00) | 1.00 (0.00) | 1.00 (0.00) | 0.98 (0.01) | 0.89 (0.30) | 0.89 (0.30) | 0.89 (0.30) | 0.98 (0.01) | 0.89 (0.30) | 0.89 (0.30) | 0.89 (0.30) | 0.59 (0.15) | 0.68 (0.14) | 0.68 (0.14) | 0.68 (0.14) | 0.83 (0.09) | 0.94 (0.04) | 0.94 (0.04) | 0.94 (0.04) |
| | **R** | 0.20 (0.09) | 0.06 (0.03) | 0.06 (0.03) | 0.06 (0.03) | 0.38 (0.06) | 0.11 (0.06) | 0.11 (0.06) | 0.11 (0.06) | 0.38 (0.06) | 0.11 (0.06) | 0.11 (0.06) | 0.11 (0.06) | 0.33 (0.06) | 0.09 (0.09) | 0.09 (0.09) | 0.09 (0.09) | 0.44 (0.10) | 0.11 (0.08) | 0.11 (0.08) | 0.11 (0.08) |
| | **F** | 0.33 (0.11) | 0.11 (0.05) | 0.11 (0.05) | 0.11 (0.05) | 0.55 (0) | 0.19 (0.10) | 0.19 (0.10) | 0.19 (0.10) | 0.55 (0) | 0.19 (0.10) | 0.19 (0.10) | 0.19 (0.10) | 0.41 (0.07) | 0.15 (0.12) | 0.15 (0.12) | 0.15 (0.12) | 0.56 (0.07) | 0.19 (0.10) | 0.19 (0.10) | 0.19 (0.10) |
| **TCI** | **T** | 0.89 (0.03) | 0.95 (0.01) | 0.95 (0.01) | 0.95 (0.01) | 0.73 (0) | 0.81 (0.03) | 0.81 (0.03) | 0.81 (0.03) | 0.73 (0) | 0.81 (0.03) | 0.81 (0.03) | 0.81 (0.03) | 0.50 (0.00) | 0.51 (0.00) | 0.51 (0.00) | 0.51 (0.00) | 0.94 (0.03) | 0.99 (0.01) | 0.99 (0.01) | 0.99 (0.01) |
| | **# >T** | 204 (87) | 57 (27) | 57 (27) | 57 (27) | 389 (67) | 110 (61) | 110 (61) | 110 (61) | 605 (222) | 148 (173) | 148 (173) | 148 (173) | 548 (180) | 119 (93) | 119 (93) | 119 (93) | 250 (114) | 70 (39) | 70 (39) | 70 (39) |
| | **# > T in EHR** | 5371 | | | | 1694 | | | | 6375 | | | | 4132 | | | | 14090 | | | |
| | **P** | 0.77 (0.16) | 0.70 (0.46) | 0.70 (0.46) | 0.70 (0.46) | 0.64 (0.06) | 0.95 (0.04) | 0.95 (0.04) | 0.95 (0.04) | 0.64 (0.06) | 0.95 (0.04) | 0.95 (0.04) | 0.95 (0.04) | 0.62 (0.15) | 0.56 (0.41) | 0.56 (0.41) | 0.56 (0.41) | 0.77 (0.20) | 0.89 (0.30) | 0.89 (0.30) | 0.89 (0.30) |
| | **R** | 0.60 (0.12) | 0.01 (0.02) | 0.01 (0.02) | 0.01 (0.02) | 0.25 (0.04) | 0.05 (0.03) | 0.05 (0.03) | 0.05 (0.03) | 0.25 (0.04) | 0.05 (0.03) | 0.05 (0.03) | 0.05 (0.03) | 0.52 (0.08) | 0.02 (0.04) | 0.02 (0.04) | 0.02 (0.04) | 0.37 (0.12) | 0.02 (0.01) | 0.02 (0.01) | 0.02 (0.01) |
| | **F** | 0.65 (0.06) | 0.03 (0.03) | 0.03 (0.03) | 0.03 (0.03) | 0.35 (0) | 0.10 (0.05) | 0.10 (0.05) | 0.10 (0.05) | 0.35 (0) | 0.10 (0.05) | 0.10 (0.05) | 0.10 (0.05) | 0.55 (0.06) | 0.03 (0.05) | 0.03 (0.05) | 0.03 (0.05) | 0.46 (0.07) | 0.04 (0.03) | 0.04 (0.03) | 0.04 (0.03) |
| **TR** | **T** | 0.83 (0.05) | 0.99 (0.01) | 0.99 (0.01) | 0.99 (0.01) | 0.83 (0) | 0.92 (0.02) | 0.92 (0.02) | 0.92 (0.02) | 0.83 (0) | 0.92 (0.02) | 0.92 (0.02) | 0.92 (0.02) | 0.50 (0.00) | 0.51 (0.00) | 0.51 (0.00) | 0.51 (0.00) | 0.93 (0.05) | 1.00 (0.00) | 1.00 (0.00) | 1.00 (0.00) |
| | **# >T** | 855 (369) | 13 (15) | 13 (15) | 13 (15) | 395 (99) | 58 (32) | 58 (32) | 58 (32) | 915 (293) | 53 (133) | 53 (133 | 53 (133 | 580 (402) | 18 (13) | 18 (13) | 18 (13) | 641 (168) | 75 (127) | 75 (127) | 75 (127) |
| | **# > T in EHR** | 42707 | | | | 42584 | | | | 43724 | | | | 23825 | | | | 102455 | | | |
| | **P** | 0.89 (0.03) | 0.99 (0.02) | 0.99 (0.02) | 0.99 (0.02) | 0.20 (0.04) | 0.16 (0.06) | 0.16 (0.06) | 0.16 (0.06) | 0.20 (0.04) | 0.16 (0.06) | 0.16 (0.06) | 0.16 (0.06) | 0.64 (0.07) | 0.77 (0.07) | 0.77 (0.07) | 0.77 (0.07) | 0.69 (0.09) | 0.91 (0.09) | 0.91 (0.09) | 0.91 (0.09) |
| **CN** | **R** | 0.47 (0.05) | 0.10 (0.11) | 0.10 (0.11) | 0.10 (0.11) | 0.18 (0.10) | 0.04 (0.05) | 0.04 (0.05) | 0.04 (0.05) | 0.18 (0.10) | 0.04 (0.05) | 0.04 (0.05) | 0.04 (0.05) | 0.39 (0.09) | 0.08 (0.09) | 0.08 (0.09) | 0.08 (0.09) | 0.27 (0.08) | 0.05 (0.04) | 0.05 (0.04) | 0.05 (0.04) |
| | **F** | 0.61 (0.04) | 0.16 (0.17) | 0.16 (0.17) | 0.16 (0.17) | 0.18 (0) | 0.06 (0.06) | 0.06 (0.06) | 0.06 (0.06) | 0.18 (0) | 0.06 (0.06) | 0.06 (0.06) | 0.06 (0.06) | 0.48 (0.06) | 0.14 (0.13) | 0.14 (0.13) | 0.14 (0.13) | 0.38 (0.07) | 0.09 (0.07) | 0.09 (0.07) | 0.09 (0.07) |

| | | | | | | | | | | | | | | | | | | | | | |
|---|---|---|---|---|---|---|---|---|---|---|---|---|---|---|---|---|---|---|---|---|---|
| | **T** | 0.91 (0.02) | 0.98 (0.02) | 0.98 (0.02) | 0.98 (0.02) | 0.81 (0) | 0.87 (0.04) | 0.87 (0.04) | 0.87 (0.04) | 0.81 (0) | 0.87 (0.04) | 0.87 (0.04) | 0.87 (0.04) | 0.51 (0.00) | 0.51 (0.00) | 0.51 (0.00) | 0.51 (0.00) | 0.97 (0.02) | 1.00 (0.00) | 1.00 (0.00) | 1.00 (0.00) |
| | **# >T** | 533 (76) | 99 (120) | 99 (120) | 99 (120) | 859 (430) | 226 (251) | 226 (251) | 226 (251) | 634 (203) | 114 (126) | 114 (126) | 114 (126) | 410 (178) | 59 (51) | 59 (51) | 59 (51) | 432 (169) | 28 (35) | 28 (35) | 28 (35) |
| | **# > T in EHR** | 68049 | | | | 67850 | | | | 68236 | | | | 53454 | | | | 129620 | | | |
| **CI** | **P** | 0.92 (0.04) | 1.00 (0.00) | 1.00 (0.00) | 1.00 (0.00) | 0.71 (0.10) | 0.96 (0.03) | 0.96 (0.03) | 0.96 (0.03) | 0.71 (0.10) | 0.96 (0.03) | 0.96 (0.03) | 0.96 (0.03) | 0.84 (0.11) | 0.98 (0.02) | 0.98 (0.02) | 0.98 (0.02) | 0.81 (0.07) | 0.86 (0.29) | 0.86 (0.29) | 0.86 (0.29) |
| | **R** | 0.74 (0.07) | 0.25 (0.12) | 0.25 (0.12) | 0.25 (0.12) | 0.75 (0.11) | 0.19 (0.11) | 0.19 (0.11) | 0.19 (0.11) | 0.75 (0.11) | 0.19 (0.11) | 0.19 (0.11) | 0.19 (0.11) | 0.71 (0.12) | 0.21 (0.18) | 0.21 (0.18) | 0.21 (0.18) | 0.67 (0.12) | 0.14 (0.12) | 0.14 (0.12) | 0.14 (0.12) |
| | **F** | 0.81 (0.03) | 0.39 (0.16) | 0.39 (0.16) | 0.39 (0.16) | 0.71 (0) | 0.31 (0.16) | 0.31 (0.16) | 0.31 (0.16) | 0.71 (0) | 0.31 (0.16) | 0.31 (0.16) | 0.31 (0.16) | 0.75 (0.05) | 0.31 (0.21) | 0.31 (0.21) | 0.31 (0.21) | 0.72 (0.05) | 0.23 (0.17) | 0.23 (0.17) | 0.23 (0.17) |
| | **T** | 0.97 (0.02) | 1.00 (0.00) | 1.00 (0.00) | 1.00 (0.00) | 0.89 (0) | 0.95 (0.01) | 0.95 (0.01) | 0.95 (0.01) | 0.89 (0) | 0.95 (0.01) | 0.95 (0.01) | 0.95 (0.01) | 0.51 (0.00) | 0.52 (0.01) | 0.52 (0.01) | 0.52 (0.01) | 0.99 (0.01) | 1.00 (0.00) | 1.00 (0.00) | 1.00 (0.00) |
| | **# >T** | 804 (121) | 251 (120) | 251 (120) | 251 (120) | 1100 (335) | 204 (116) | 204 (116) | 204 (116) | 877 (287) | 213 (189) | 213 (189) | 213 (189) | 838 (223) | 150 (136) | 150 (136) | 150 (136) | 834 (198) | 126 (165) | 126 (165) | 126 (165) |
| | **# > T in EHR** | 105878 | | | | 177395 | | | | 99574 | | | | 98883 | | | | 166550 | | | |
| **CC** | **P** | 0.82 (0.09) | 0.97 (0.03) | 0.97 (0.03) | 0.97 (0.03) | 0.69 (0.14) | 0.99 (0.03) | 0.99 (0.03) | 0.99 (0.03) | 0.69 (0.14) | 0.99 (0.03) | 0.99 (0.03) | 0.99 (0.03) | 0.82 (0.08) | 0.97 (0.03) | 0.97 (0.03) | 0.97 (0.03) | 0.78 (0.09) | 0.95 (0.02) | 0.95 (0.02) | 0.95 (0.02) |
| | **R** | 0.83 (0.07) | 0.49 (0.21) | 0.49 (0.21) | 0.49 (0.21) | 0.83 (0.12) | 0.14 (0.17) | 0.14 (0.17) | 0.14 (0.17) | 0.83 (0.12) | 0.14 (0.17) | 0.14 (0.17) | 0.14 (0.17) | 0.80 (0.09) | 0.46 (0.17) | 0.46 (0.17) | 0.46 (0.17) | 0.78 (0.06) | 0.32 (0.19) | 0.32 (0.19) | 0.32 (0.19) |
| | **F** | 0.82 (0.02) | 0.61 (0.23) | 0.61 (0.23) | 0.61 (0.23) | 0.73 (0) | 0.22 (0.20) | 0.22 (0.20) | 0.22 (0.20) | 0.73 (0) | 0.22 (0.20) | 0.22 (0.20) | 0.22 (0.20) | 0.80 (0.03) | 0.60 (0.17) | 0.60 (0.17) | 0.60 (0.17) | 0.77 (0.03) | 0.44 (0.23) | 0.44 (0.23) | 0.44 (0.23) |
| | **T** | 0.96 (0.03) | 1.00 (0.00) | 1.00 (0.00) | 1.00 (0.00) | 0.88 (0) | 0.96 (0.01) | 0.96 (0.01) | 0.96 (0.01) | 0.88 (0) | 0.96 (0.01) | 0.96 (0.01) | 0.96 (0.01) | 0.51 (0.00) | 0.52 (0.01) | 0.52 (0.01) | 0.52 (0.01) | 0.99 (0.01) | 1.00 (0.00) | 1.00 (0.00) | 1.00 (0.00) |
| | **# >T** | 1044 (212) | 506 (232) | 506 (232) | 506 (232) | 1287 (458) | 151 (184) | 151 (184) | 151 (184) | 996 (197) | 476 (191) | 476 (191) | 476 (191) | 1026 (228) | 336 (206) | 336 (206) | 336 (206) | 906 (167) | 161 (114) | 161 (114) | 161 (114) |
| | **# > T in EHR** | 123519 | | | | 174975 | | | | 115115 | | | | 114172 | | | | 171534 | | | |
| **CCI** | **P** | 0.99 (0.01) | 1.00 (0.00) | 1.00 (0.00) | 1.00 (0.00) | 0.98 (0.01) | 0.99 (0.01) | 0.99 (0.01) | 0.99 (0.01) | 0.98 (0.01) | 0.99 (0.01) | 0.99 (0.01) | 0.99 (0.01) | 0.39 (0.07) | 0.72 (0.14) | 0.72 (0.14) | 0.72 (0.14) | 0.70 (0.10) | 0.94 (0.04) | 0.94 (0.04) | 0.94 (0.04) |

| | | | | | | | | | | | | | | | | | | | | | |
|---|---|---|---|---|---|---|---|---|---|---|---|---|---|---|---|---|---|---|---|---|---|
| | **R** | 0.35 (0.11) | 0.06 (0.04) | 0.06 (0.04) | 0.06 (0.04) | 0.49 (0.12) | 0.14 (0.12) | 0.14 (0.12) | 0.14 (0.12) | 0.49 (0.12) | 0.14 (0.12) | 0.14 (0.12) | 0.14 (0.12) | 0.50 (0.12) | 0.04 (0.01) | 0.04 (0.01) | 0.04 (0.01) | 0.35 (0.13) | 0.08 (0.05) | 0.08 (0.05) | 0.08 (0.05) |
| | **F** | 0.50 (0.13) | 0.12 (0.07) | 0.12 (0.07) | 0.12 (0.07) | 0.64 (0) | 0.23 (0.17) | 0.23 (0.17) | 0.23 (0.17) | 0.64 (0) | 0.23 (0.17) | 0.23 (0.17) | 0.23 (0.17) | 0.43 (0.08) | 0.08 (0.02) | 0.08 (0.02) | 0.08 (0.02) | 0.44 (0.09) | 0.14 (0.08) | 0.14 (0.08) | 0.14 (0.08) |
| | **T** | 0.76 (0.03) | 0.87 (0.03) | 0.87 (0.03) | 0.87 (0.03) | 0.65 (0) | 0.74 (0.04) | 0.74 (0.04) | 0.74 (0.04) | 0.65 (0) | 0.74 (0.04) | 0.74 (0.04) | 0.74 (0.04) | 0.50 (0.00) | 0.51 (0.00) | 0.51 (0.00) | 0.51 (0.00) | 0.89 (0.04) | 0.98 (0.01) | 0.98 (0.01) | 0.98 (0.01) |
| | **# >T** | 349 (109) | 63 (40) | 63 (40) | 63 (40) | 497 (130) | 142 (123) | 142 (123) | 142 (123) | 1271 (266) | 60 (21) | 60 (21) | 60 (21) | 538 (266) | 82 (53) | 82 (53) | 82 (53) | 399 (119) | 79 (73) | 79 (73) | 79 (73) |
| | **# > T in EHR** | 2284 | | | | 161 | | | | 7461 | | | | 1391 | | | | 8945 | | | |
| **CR** | **P** | 0.67 (0.20) | 0.89 (0.30) | 0.89 (0.30) | 0.89 (0.30) | 0.45 (0.06) | 0.83 (0.07) | 0.83 (0.07) | 0.83 (0.07) | 0.45 (0.06) | 0.83 (0.07) | 0.83 (0.07) | 0.83 (0.07) | 0.52 (0.07) | 0.50 (0.13) | 0.50 (0.13) | 0.50 (0.13) | 0.65 (0.14) | 0.78 (0.39) | 0.78 (0.39) | 0.78 (0.39) |
| | **R** | 0.60 (0.13) | 0.09 (0.06) | 0.09 (0.06) | 0.09 (0.06) | 0.27 (0.03) | 0.08 (0.04) | 0.08 (0.04) | 0.08 (0.04) | 0.27 (0.03) | 0.08 (0.04) | 0.08 (0.04) | 0.08 (0.04) | 0.54 (0.08) | 0.12 (0.19) | 0.12 (0.19) | 0.12 (0.19) | 0.42 (0.09) | 0.03 (0.05) | 0.03 (0.05) | 0.03 (0.05) |
| | **F** | 0.59 (0.08) | 0.16 (0.11) | 0.16 (0.11) | 0.16 (0.11) | 0.34 (0) | 0.14 (0.06) | 0.14 (0.06) | 0.14 (0.06) | 0.34 (0) | 0.14 (0.06) | 0.14 (0.06) | 0.14 (0.06) | 0.52 (0.04) | 0.13 (0.16) | 0.13 (0.16) | 0.13 (0.16) | 0.49 (0.06) | 0.06 (0.08) | 0.06 (0.08) | 0.06 (0.08) |
| | **T** | 0.81 (0.06) | 0.97 (0.02) | 0.97 (0.02) | 0.97 (0.02) | 0.80 (0) | 0.90 (0.02) | 0.90 (0.02) | 0.90 (0.02) | 0.80 (0) | 0.90 (0.02) | 0.90 (0.02) | 0.90 (0.02) | 0.50 (0.00) | 0.51 (0.00) | 0.51 (0.00) | 0.51 (0.00) | 0.91 (0.05) | 0.99 (0.01) | 0.99 (0.01) | 0.99 (0.01) |
| | **# >T** | 1104 (765) | 93 (65) | 93 (65) | 93 (65) | 623 (136) | 94 (49) | 94 (49) | 94 (49) | 1081 (331) | 259 (423) | 259 (423) | 259 (423) | 717 (327) | 37 (55) | 37 (55) | 37 (55) | 2424 (3078) | 84 (129) | 84 (129) | 84 (129) |
| | **# > T in EHR** | 55359 | | | | 48311 | | | | 48898 | | | | 38868 | | | | 124475 | | | |

**Supplementary Table 6:** Precision (P), Recall (R) Maximum F score (FB1, FB1/2, FB1/4, FB1/8), probability threshold (T) at four F scores, number of patients in test set above threshold (\#>T), and number of patients in EHR above max F1 threshold (\#>T in EHR). Standard deviation in (). See Method “Model” and Supplementary Table 1 for case-control, model, and evaluator abbreviations' definitions**.**

| Model | Sensitivity | Specificity | Positive Predictive Value | Negative Predictive Value |
|---|---|---|---|---|
| **LR** | 0.256 (0.247,0.265) | 0.997 (0.997,0.997) | 5.92e-4 (5.71e-4,6.14e-4) | 0.998 (0.998,0.998) |
| **EN** | 0.192 (0.186,0.197) | 0.997 (0.996,0.997) | 4.43e-4 (4.31e-4, 4.56e-4) | 0.998 (0.998,0.998) |
| **RF** | 0.255 (0.247,0.264) | 0.997 (0.997,0.997) | 5.90e-4 (5.71e-4, 6.10e-4) | 0.998 (0.998, 0.998) |
| **AB** | 0.388 (0.370, 0.406) | 0.998 (0.998,0.998) | 8.95e-4 (8.55e-4, 9.37e-4) | 0.998 (0.998,0.998) |
| **GB** | 0.286 (0.274, 0.297) | 0.996 (0.995,0.996) | 6.61e-4 (6.35e-4,6.88 e-4) | 0.998 (0.998,0.998) |

**Supplementary Table 7:** Sensitivity, specificity, positive predictive value, and negative predictive value of models on holdout test set. Standard deviation in (). See Supplementary Table 1 for case-control, model, and evaluator abbreviations' definitions.

#### SUPPLEMENTARY FIGURES

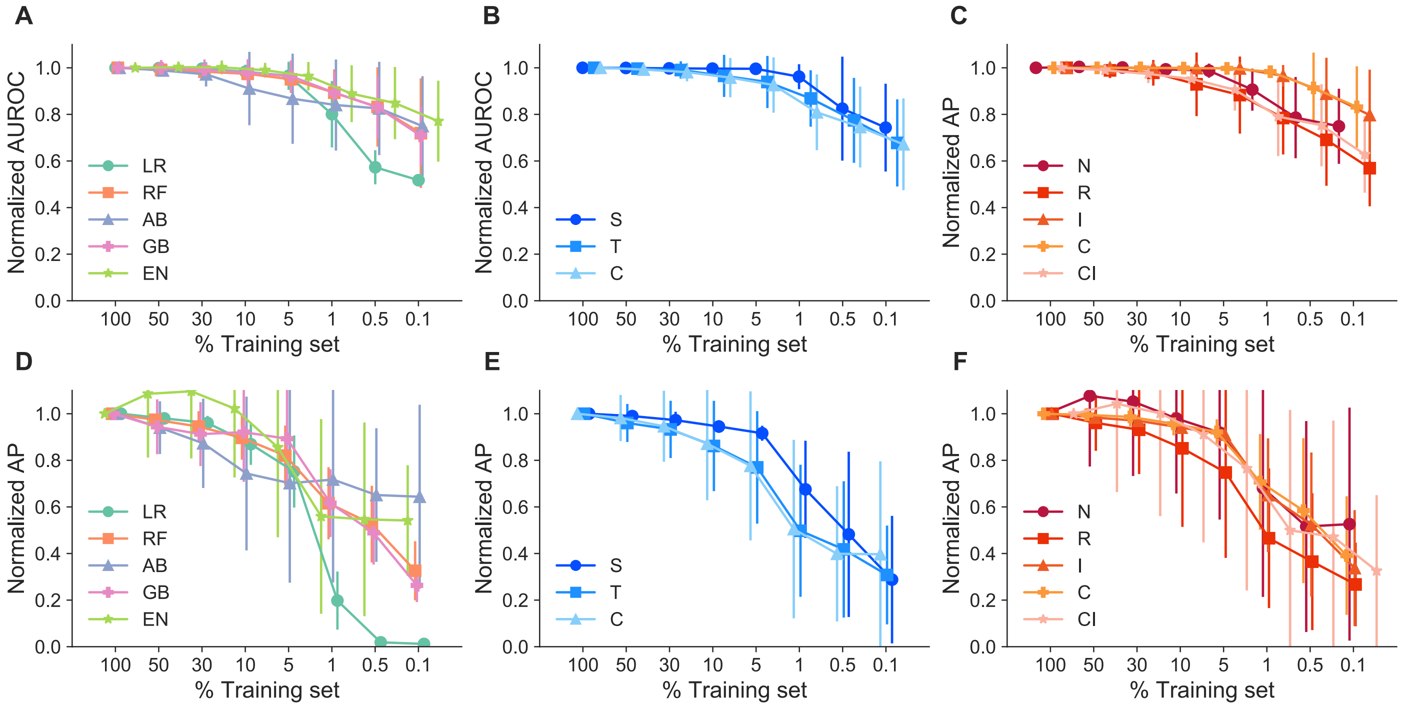


**Supplementary Figure 1:** Robustness of models. Top: Normalized area under the receiver operating curve (AUROC) across all case-control combinations, **A:** stratified by classifier type (LR, RF, AB, GB, and EN), **B:** stratified by case type (first letter S,T,C), **C:** stratified by control type (second and third letters: N, R, I,C,CI). Bottom: Normalized Average precision score (AP) values across all case-control combinations, **D:** stratified by classifier type (LR, RF, AB, GB, and EN, **E:** stratified by case type (first letter S,T,C), **F:** stratified by control type (second and third letters: N, R, I,C,CI). Neural Network Models and EN GI model were removed from normalized graphs due to high variance and obfuscation of other models' trends.

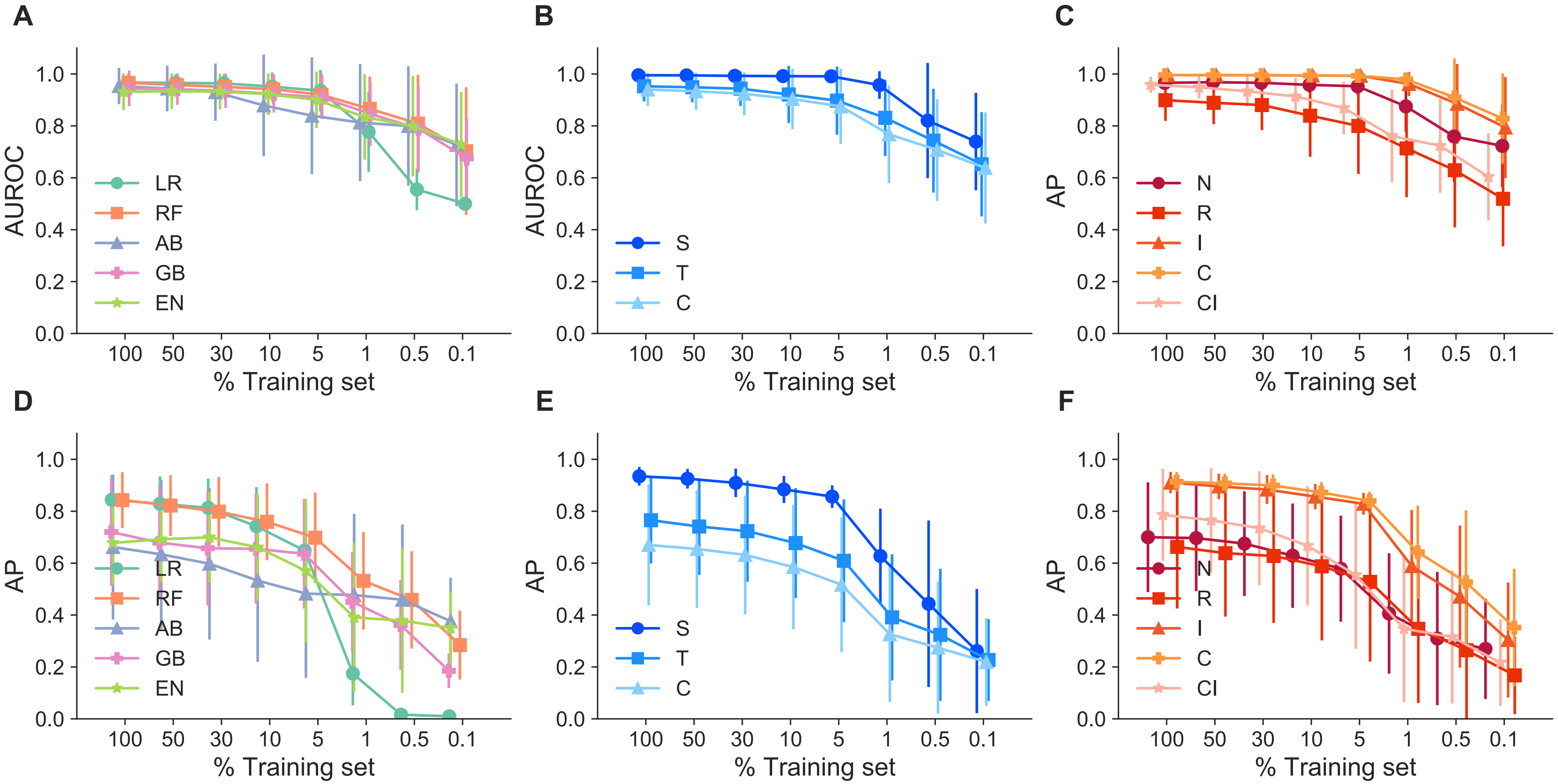


**Supplementary Figure 2:** Robustness of models. Top panel, Left to Right: Area under the receiver operating curve (AUROC) across all case-control combinations, A: stratified by classifier type (LR, RF, AB, GB, and EN), B: AUROC across all models stratified by case type (first letter: S, T, C), C: AUROC across all case-control combinations, stratified by control type (second and third letters: N, R, I,C,CI). Bottom Panel, left to right:  D: Average precision score (AP) values across all case-control combinations, stratified by model, E: AP across all models stratified by case type, F AP across all models stratified by control type. Neural Network Models and EN GI model were removed from normalized graphs due to high variance and obfuscation of other models' trends.

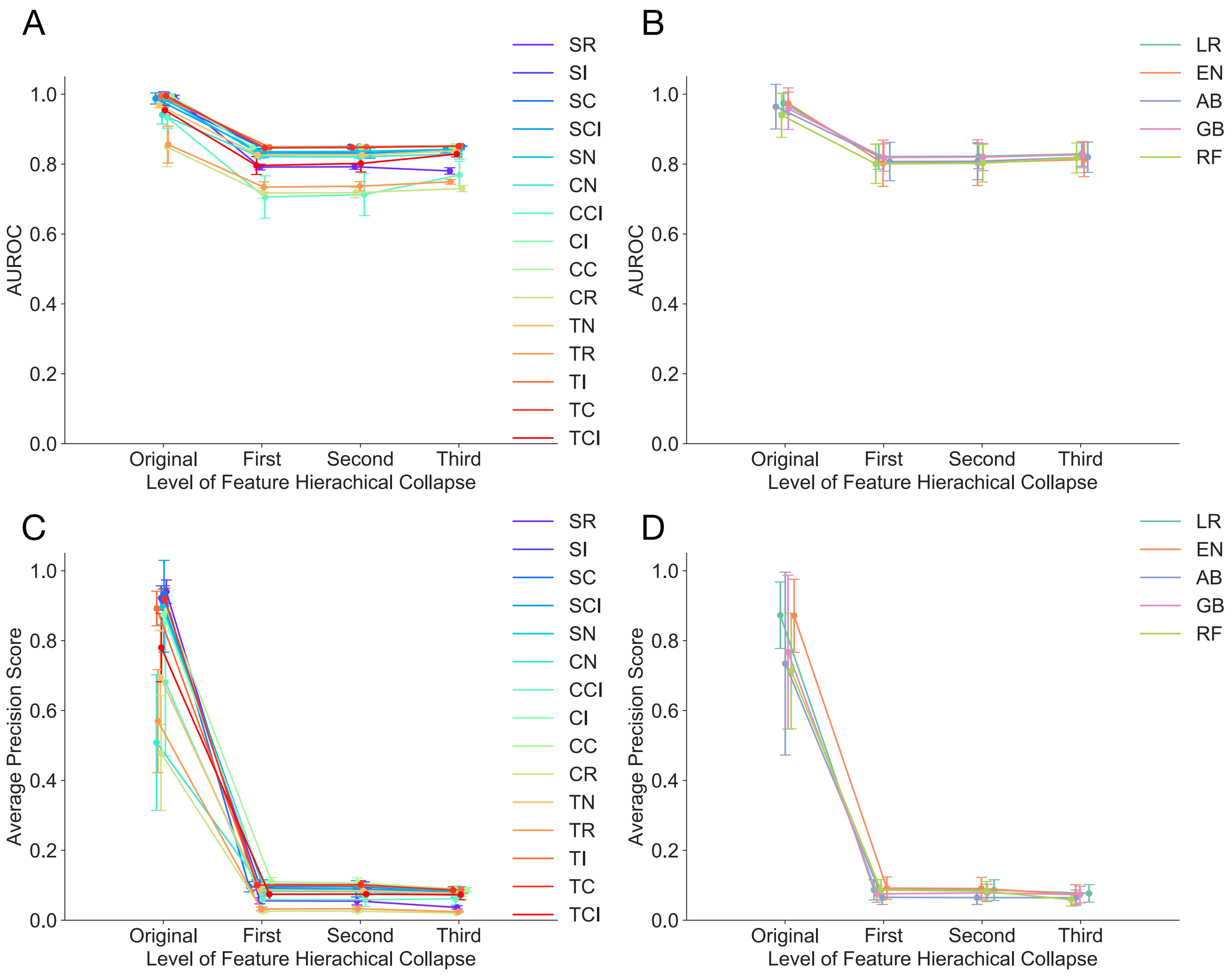

**Supplementary Figure 3:** Performance of models at increasing level of feature hierarchical collapse. Logistic regression with L1 penalty (LR), Logistic Regression with elastic net penalty (EN), random forest (RF), AdaBoost (AB), gradient boosting (GB), and a two-layer neural network (NN). Top panel, Left to Right: A shows area under the receiver operating curve AUROC stratified by case-control type, B shows AUROC stratified by classifier type versus the level of abstraction with `First' being the lowest level of abstraction and `Third' being the highest. Bottom panel, Left to Right: C shows average precision score (AP) stratified by case-control type, D shows AP stratified by classifier type versus the level of abstract with `First' being the lowest level of abstraction and `Third' being the highest. See Supplementary Table 1 for case-control abbreviations' definitions.

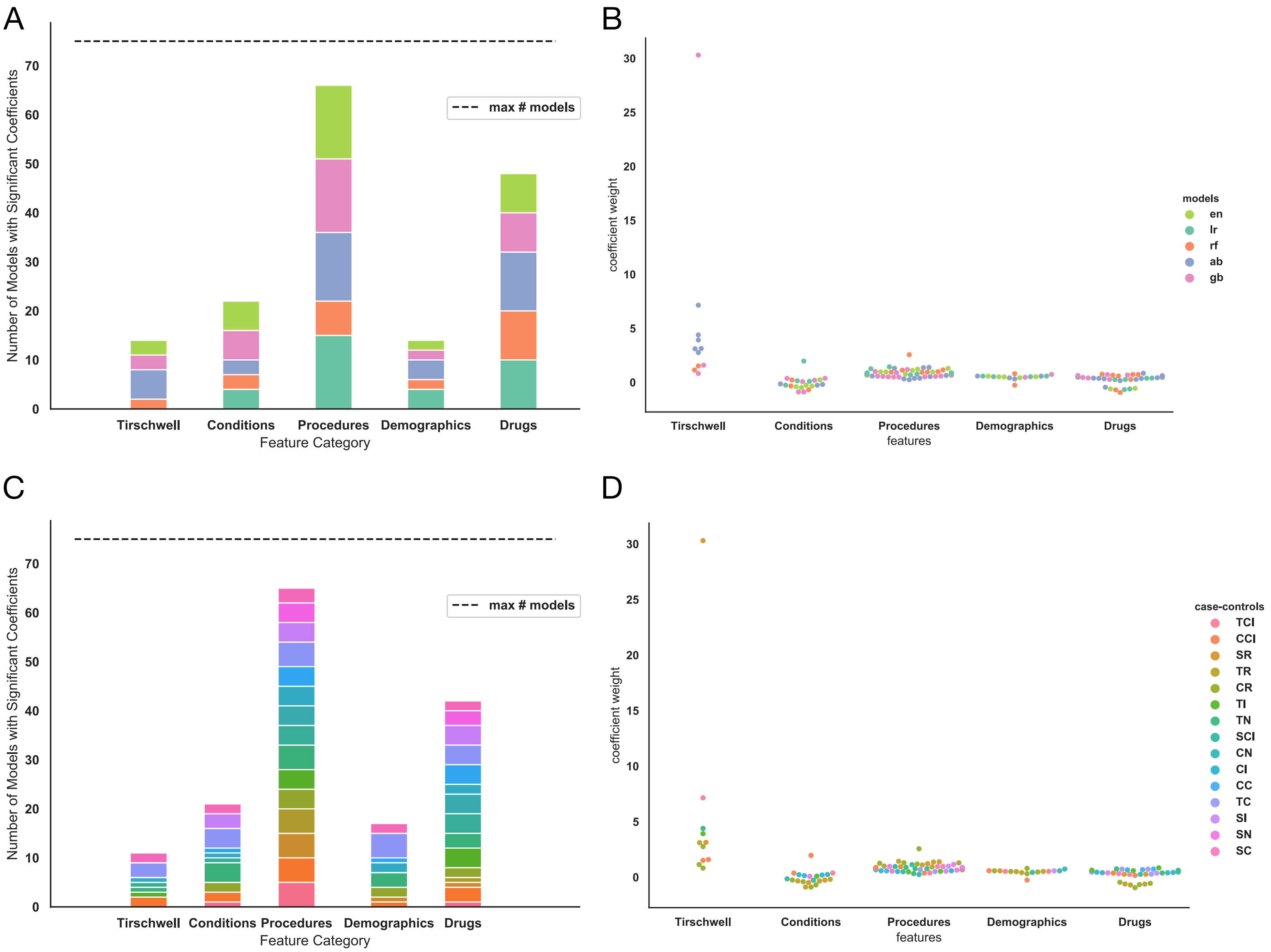
A
B
C
D
max # models
Number of Models with Significant Coefficients
Feature Category
Tirschwell
Conditions
Procedures
Demographics
Drugs
0
10
20
30
40
50
60
70
coefficient weight
features
5
15
25
models
en
lr
rf
ab
gb
case-controls
TCI
CCI
SR
TR
CR
TI
TN
SCI
CN
CI
CC
TC
SI
SN
SC

**Supplementary Figure 4**: Feature category contribution to model fits. Top panel Left to right: A shows the total number of models with a significant coefficient weight for each feature category, stratified by classifier type. B shows the value of the significant coefficient weights, stratified by classifier type. Bottom panel Left to right: C shows the total number of models with a significant coefficient weight for each feature category, stratified by case-control type. D shows the value of the significant coefficient weights, stratified by case-control type. TCI Random Forest model and CCI EN model had significant coefficients in all categories, and all of these weights were greater than 1 x $10^{14}$, so they would be plotted beyond the scope of the y axis in B and D.

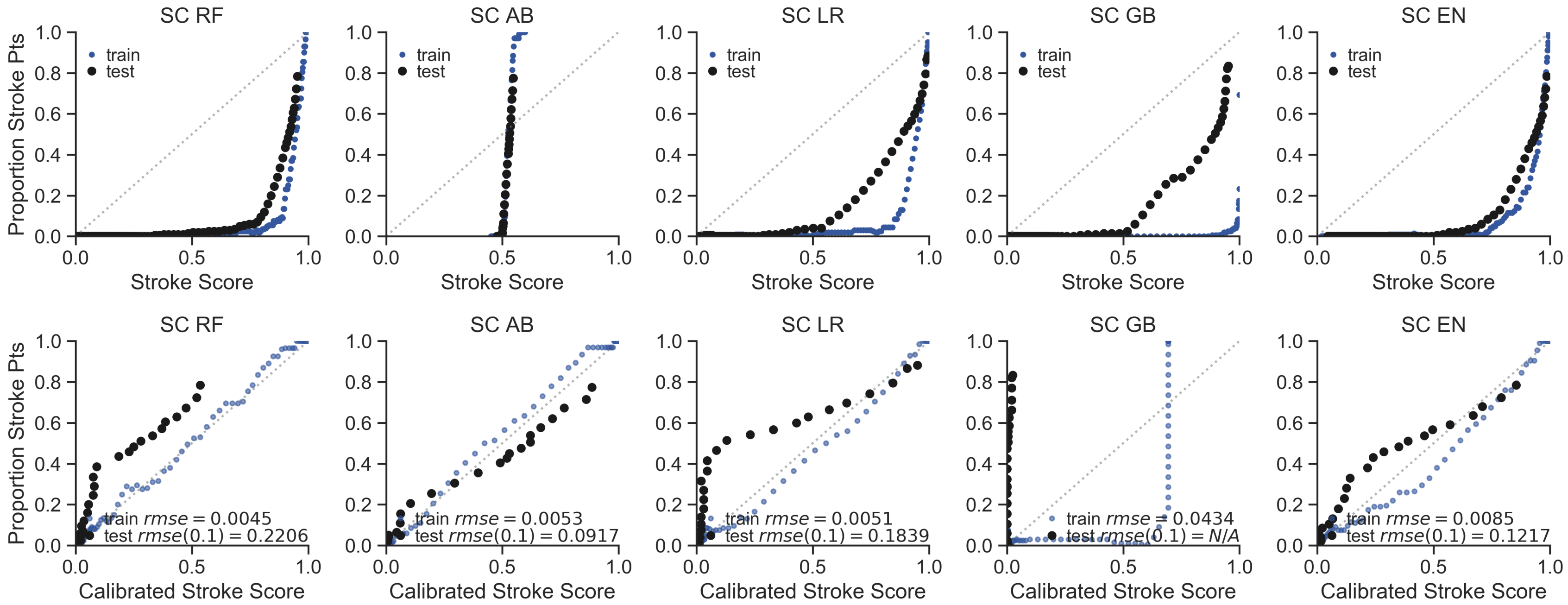


**Supplementary Figure 5:** Classifier type with stroke service cases and without cerebrovascular disease (SC) case-control combination varies in calibration success between stroke score, or model probability, and actual proportion of patients at each probability. Top panel plots the proportion of stroke case patients within a bin of 100 patients with similar stroke scores for models trained with stroke service cases and controls without cerebrovascular disease ICD9 or 10 codes. The black circles plot the mean test stroke score across all training set folds, while the blue circles plot the training fold stroke scores combined. Classifier type varies from left to right: Random Forest (RF), AdaBoost (AB), Logistic Regression with L1 penalty (LR), Gradient Boosting (GB), Logistic regression with elastic net penalty (EN). Bottom panel plots the proportion of stroke patients within a bin of 100 test set patients with similar stroke scores versus the calibrated test scores at specific scores rounded to the nearest thousandth (Black dots). The test scores are calibrated from an empirical distribution determined from the training set (See Supplementary Methods). The blue dots plot the proportion of stroke patients within a bin of 100 bootstrapped training set patients with similar stroke scores versus the calibrated training set scores rounded to the nearest thousandth. The light grey dots show perfect calibration.

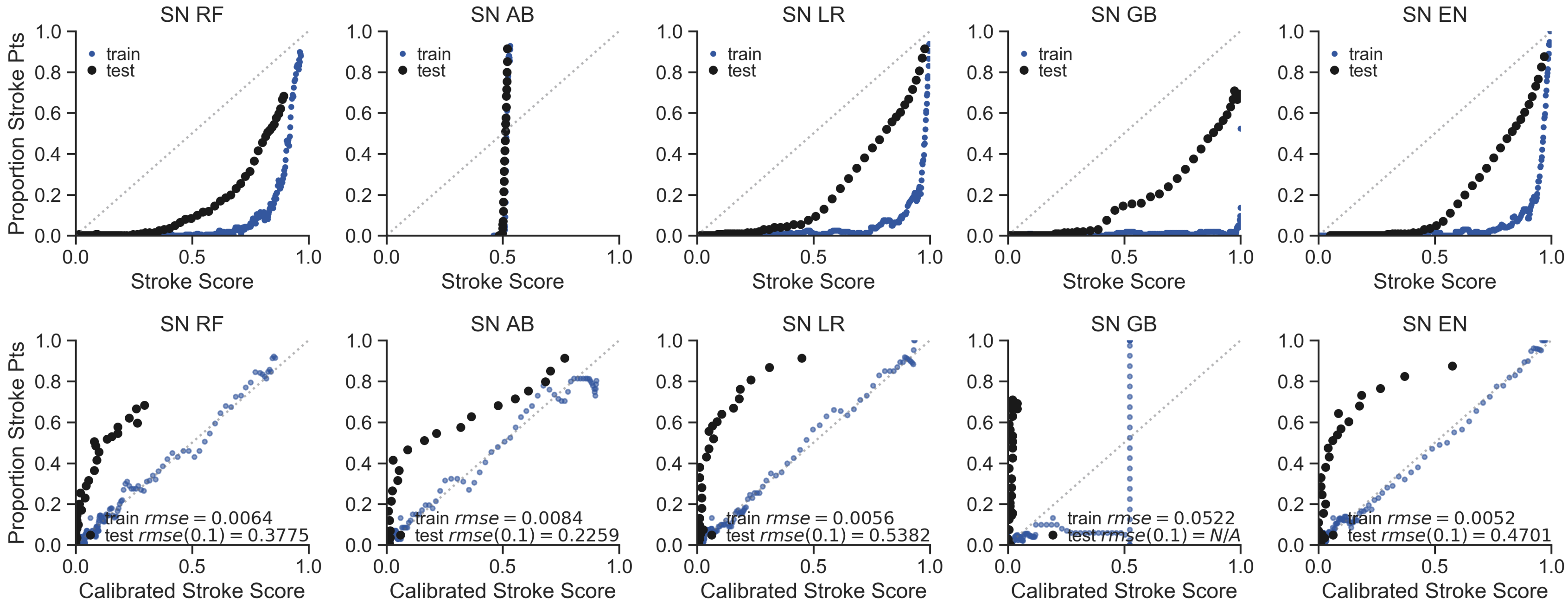


**Supplementary Figure 6:** Classifier type with Stroke Service Cases, Stroke Mimetic Controls (SN) case-control combination varies in calibration success between stroke score and actual proportion of patients at each probability.

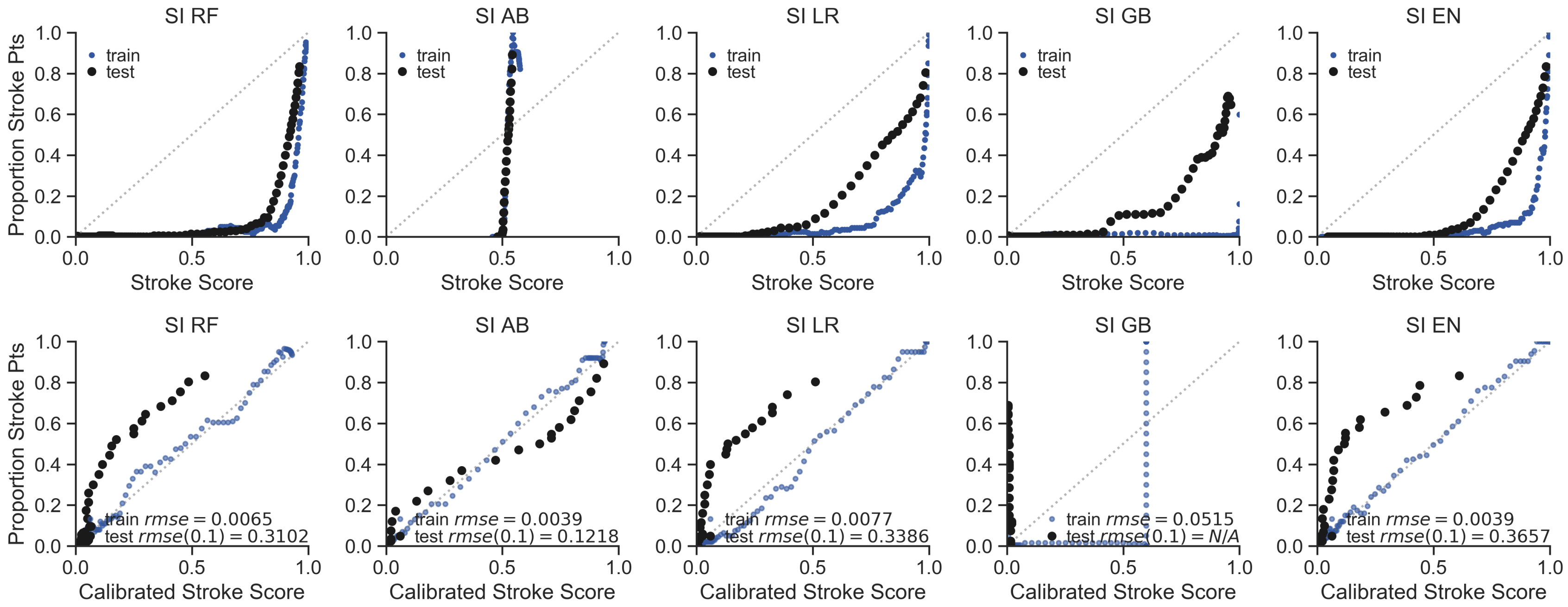


**Supplementary Figure 7:** Classifier type with Stroke Service Cases, Controls without T-L codes for AIS (SI) case-control combination varies in calibration success between stroke score and actual proportion of patients at each probability.

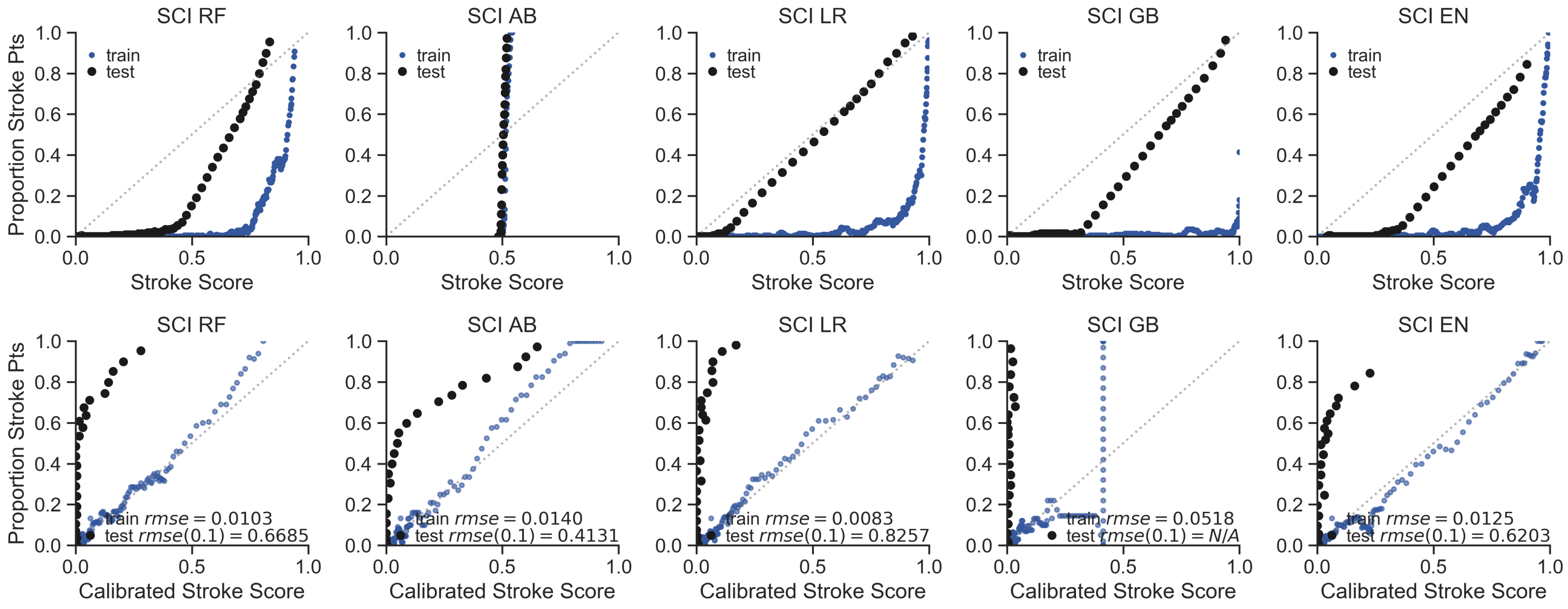


**Supplementary Figure 8:** Classifier type with Stroke Service Cases, Controls with ICD9 or ICD10 codes for CvD, and without T-L codes for AIS (SCI) case-control combination varies in calibration success between stroke score and actual proportion of patients at each probability.

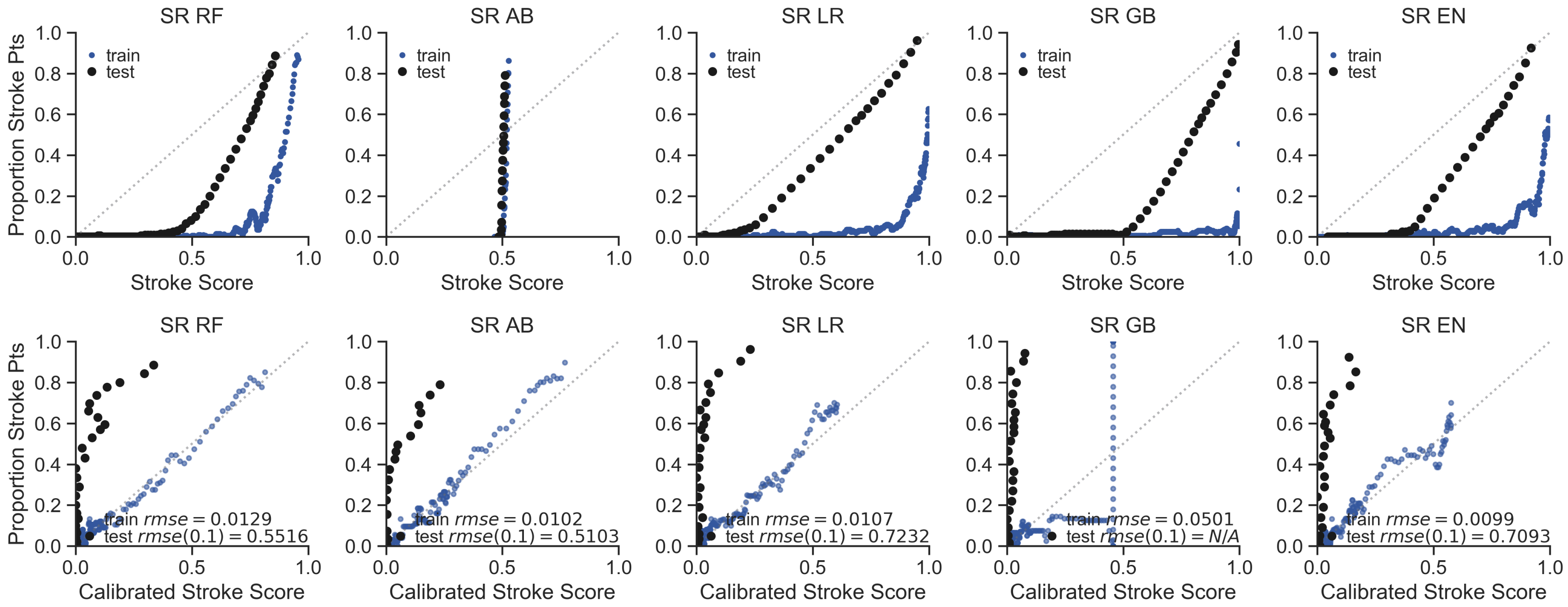


**Supplementary Figure 9:** Classifier type with Stroke Service Cases, Random patients in the EHR as controls (SR) case-control combination varies in calibration success between stroke score and actual proportion of patients at each probability.

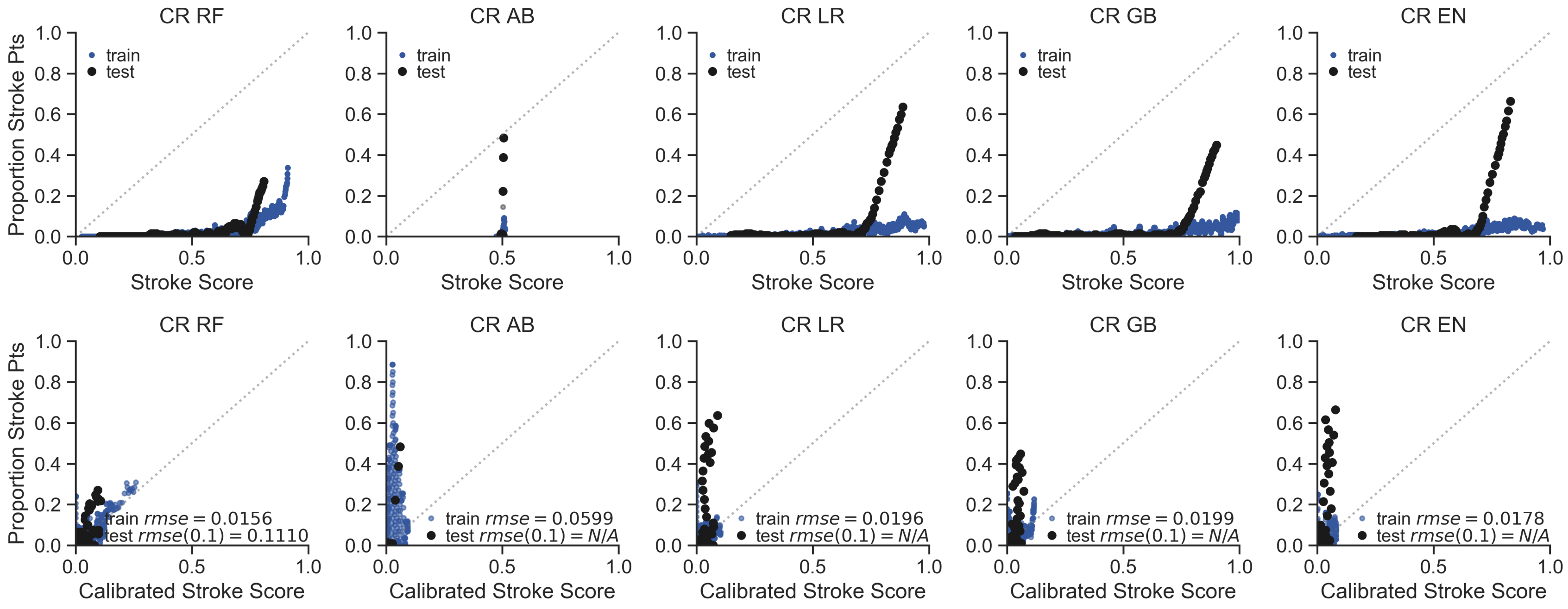


**Supplementary Figure 10:** Classifier type with Cases with ICD9 or ICD10 codes for  CvD, Random patients in the EHR as controls (CR) case-control combination varies in calibration success between stroke score and actual proportion of patients at each probability.

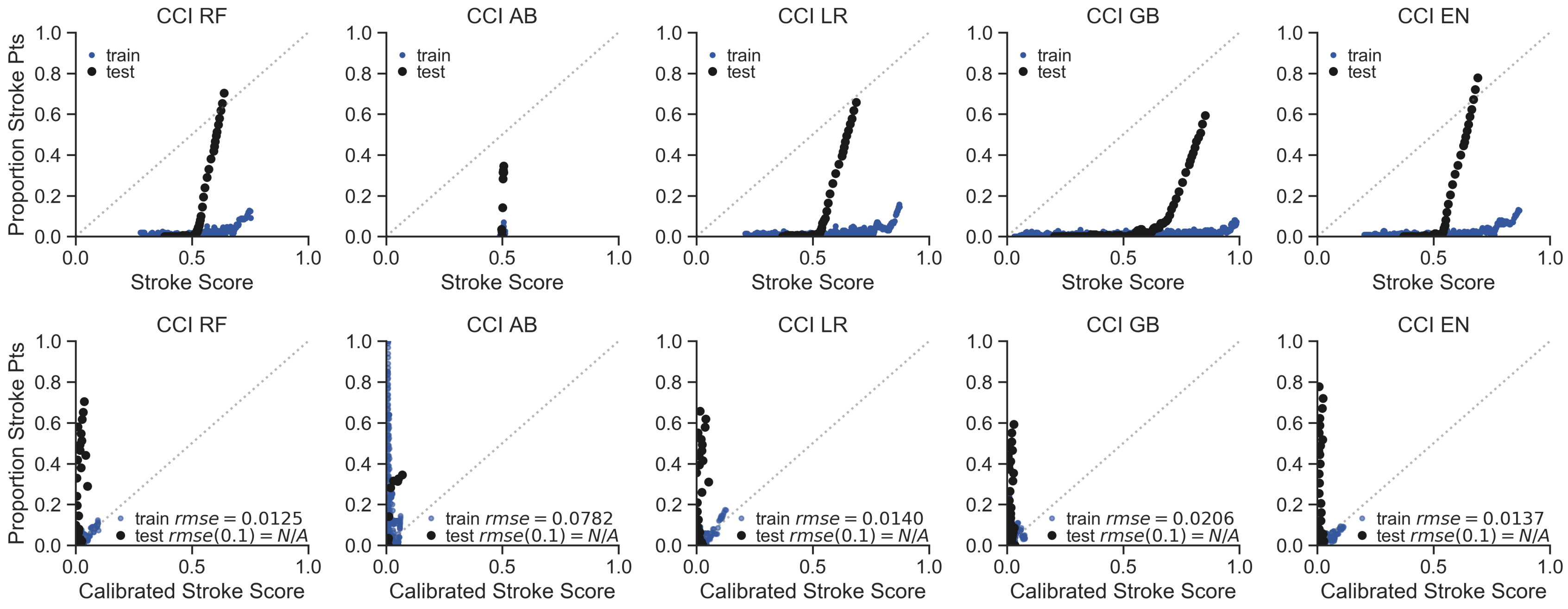


**Supplementary Figure 11:** Classifier type with Cases with ICD9 or ICD10 codes for  CvD, Controls with ICD9 or ICD10 codes for CvD and without T-L codes for AIS (CCI) case-control combination varies in calibration success between stroke score and actual proportion of patients at each probability.

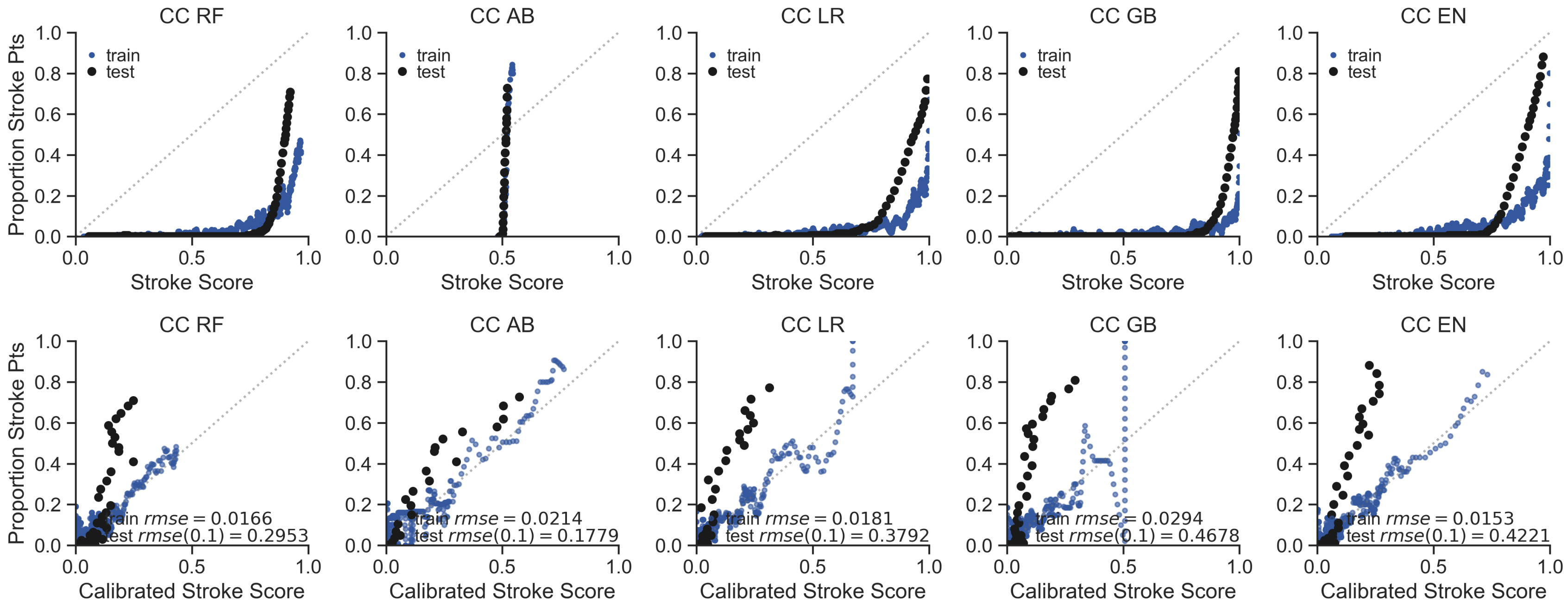


**Supplementary Figure 12:** Classifier type with Cases with ICD9 or ICD10 codes for CvD, Controls without ICD9 or ICD10 codes for CvD (CC) case-control combination varies in calibration success between stroke score and actual proportion of patients at each probability.

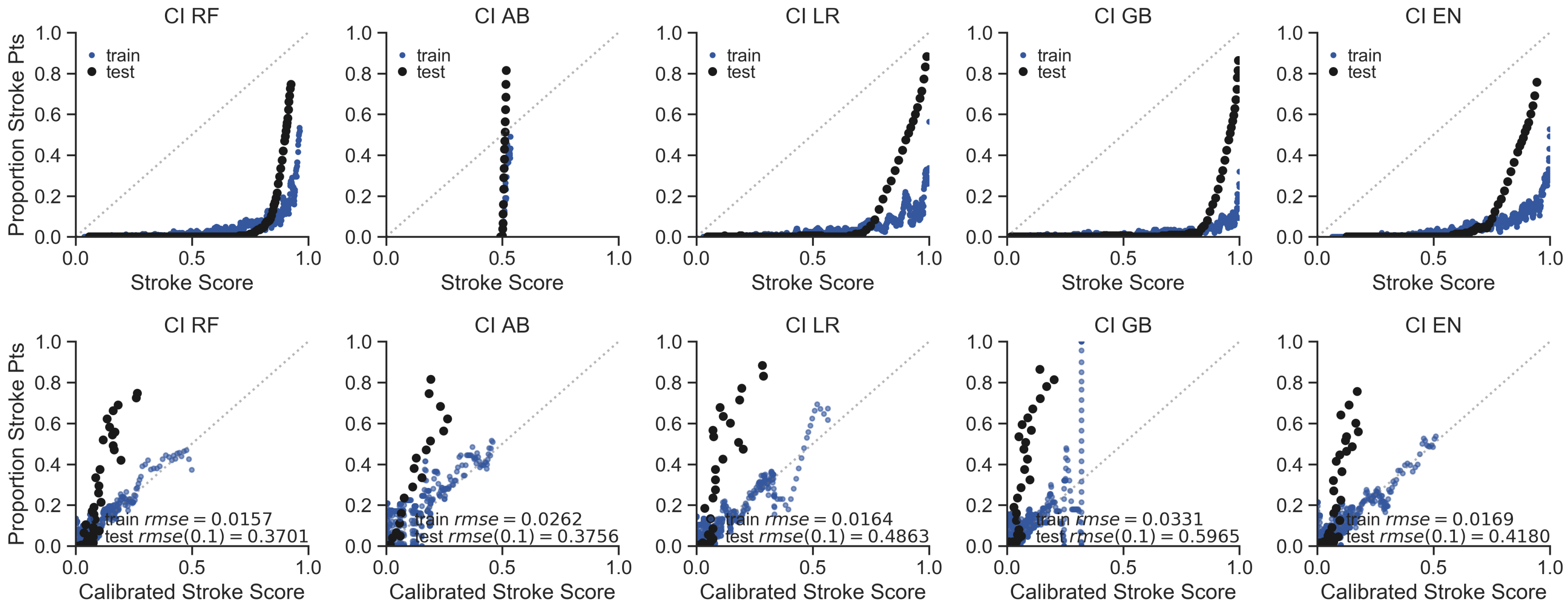


**Supplementary Figure 13:** Classifier type with Cases with ICD9 or ICD10 codes for CvD, Controls without T-L codes for AIS (CI) case-control combination varies in calibration success between stroke score and actual proportion of patients at each probability.

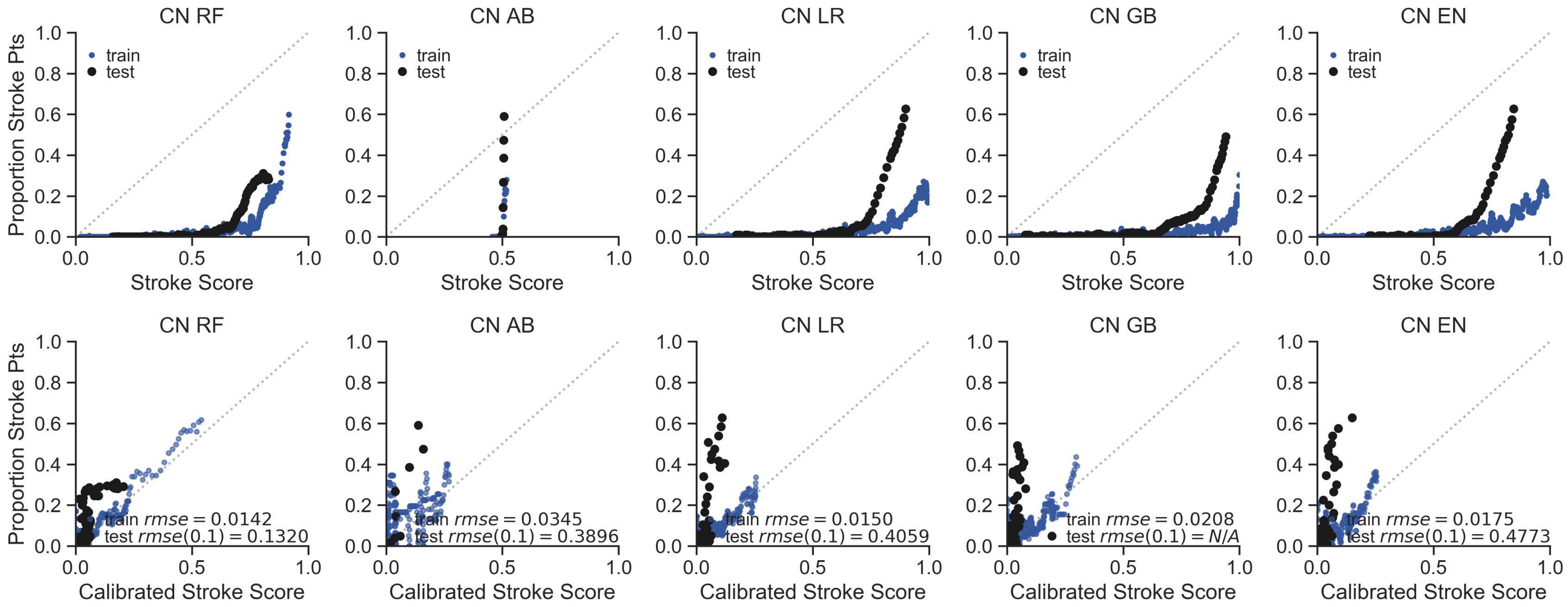


**Supplementary Figure 14:** Classifier type with Cases with ICD9 or ICD10 codes for CvD, Stroke Mimetic Controls (CN) case-control combination varies in calibration success between stroke score and actual proportion of patients at each probability.

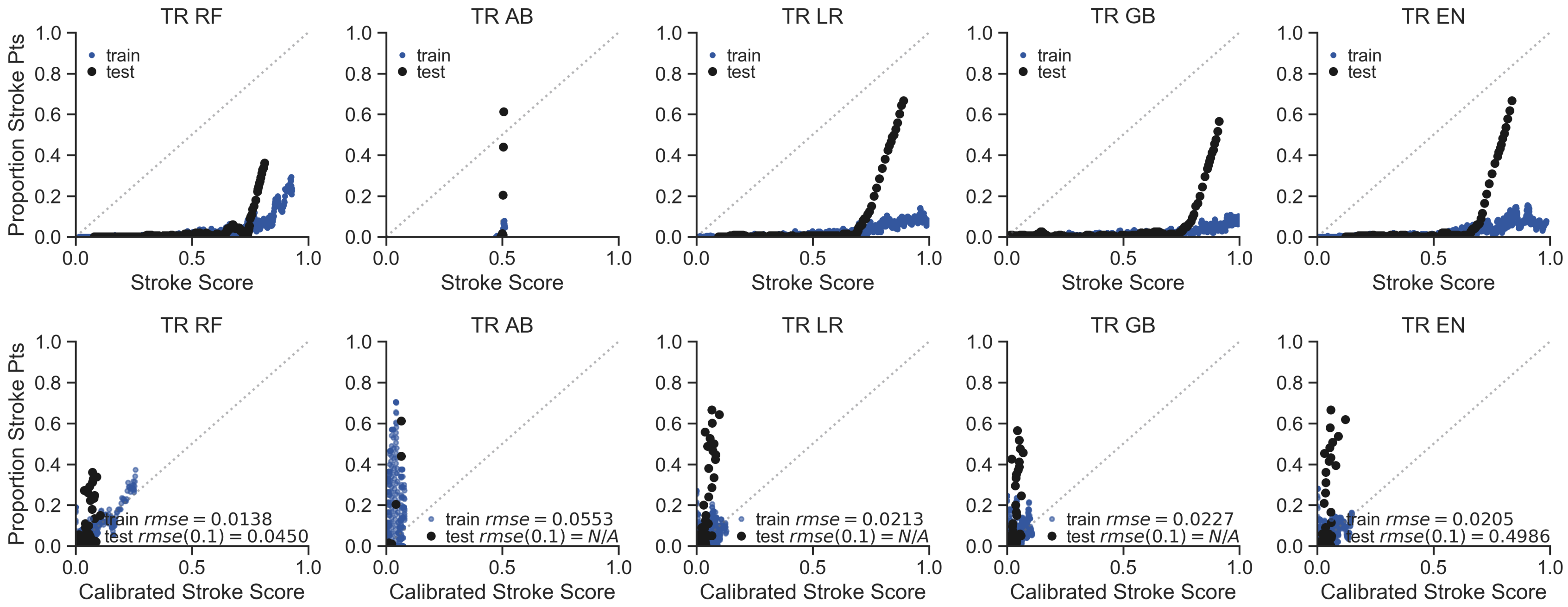


**Supplementary Figure 15:** Classifier type with Cases with T-L codes for AIS, Random patients in the EHR as controls (TR) case-control combination varies in calibration success between stroke score and actual proportion of patients at each probability.

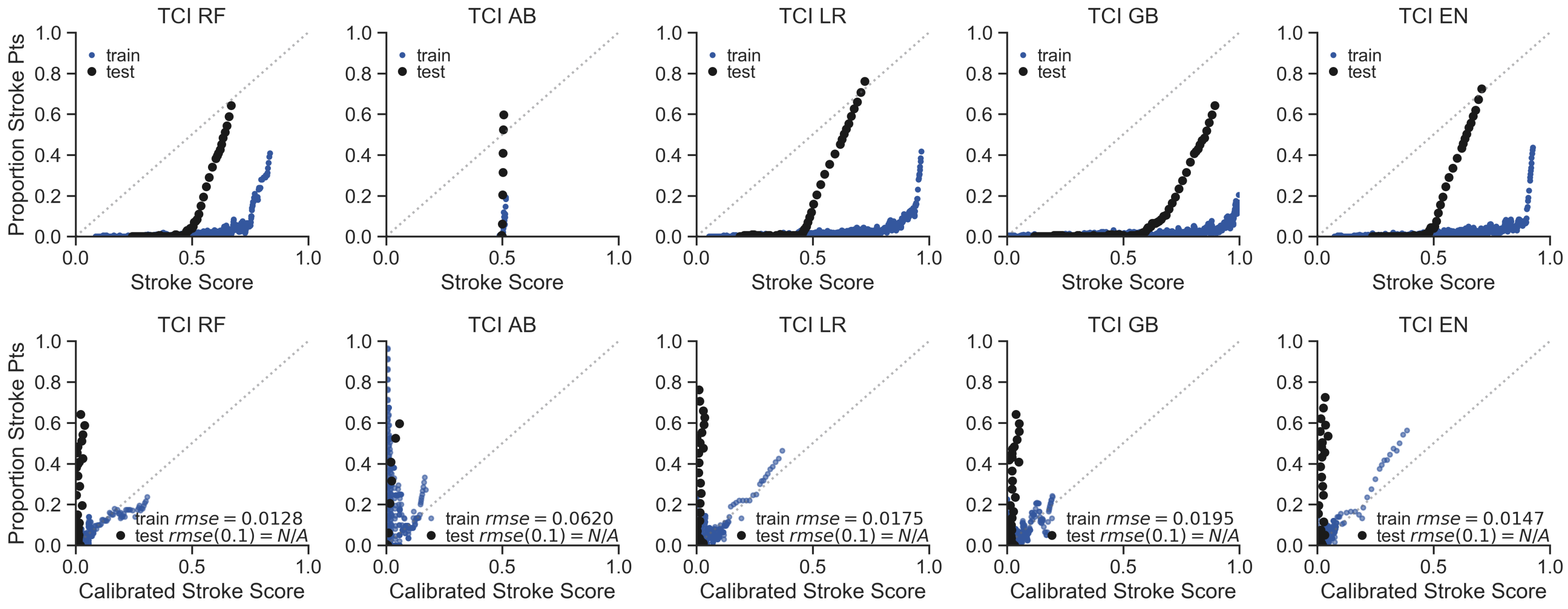


**Supplementary Figure 16:** Classifier type with Cases with T-L codes for AIS, Controls with ICD9 or ICD10 codes for CvD, and without T-L codes for AIS (TCI) case-control combination varies in calibration success between stroke score and actual proportion of patients at each probability.

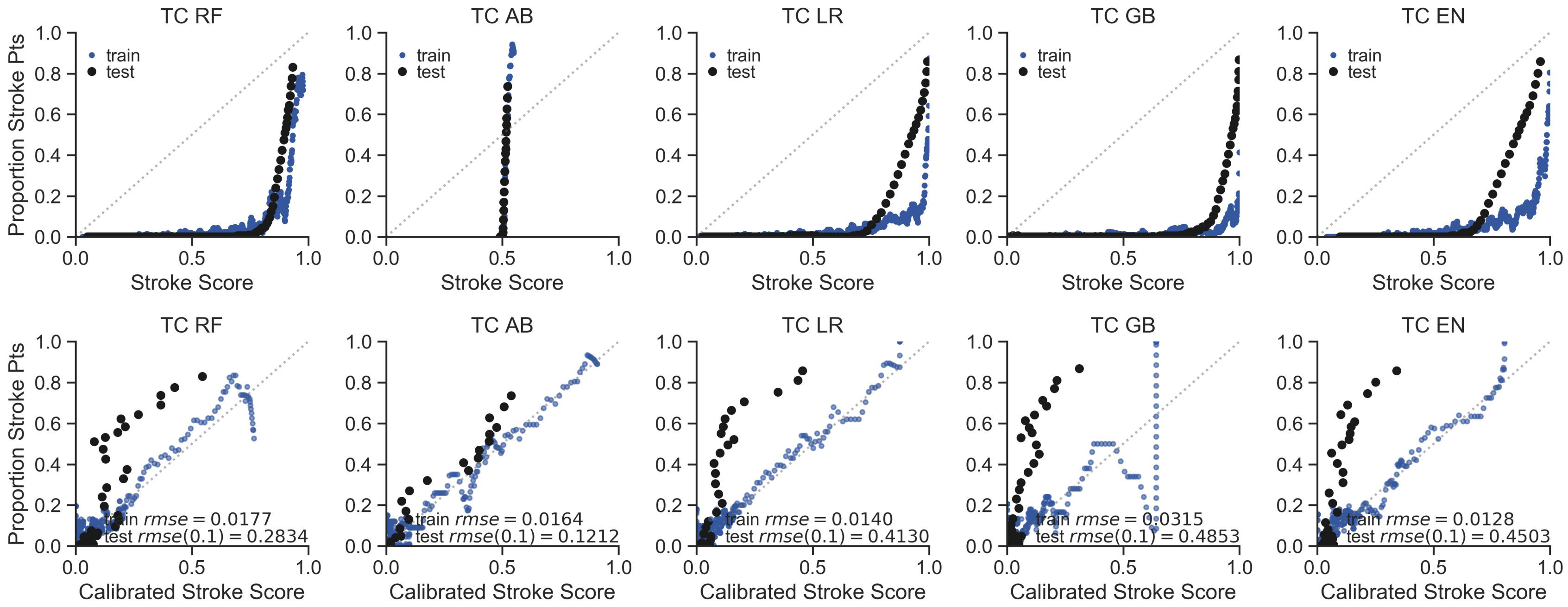


**Supplementary Figure 17:** Classifier type with Cases with T-L codes for AIS, Controls without ICD9 or ICD10 codes for CvD (TC) case-control combination varies in calibration success between stroke score and actual proportion of patients at each probability.

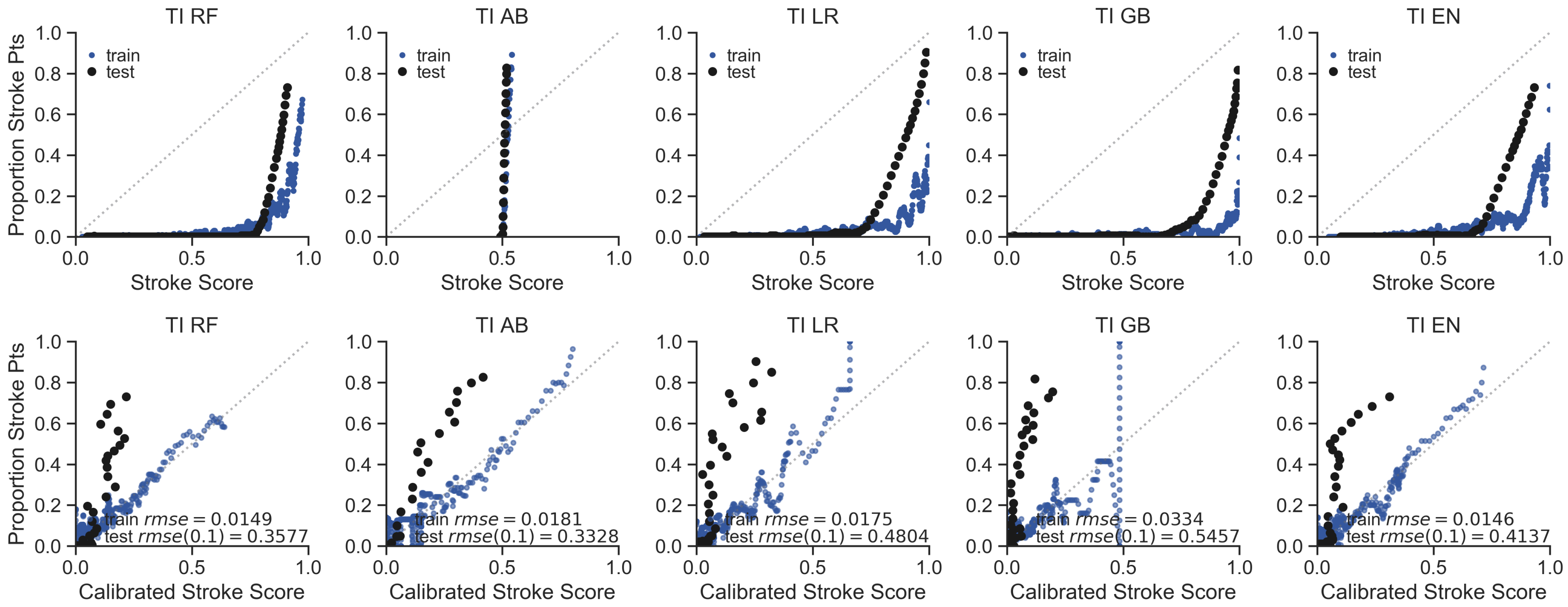


**Supplementary Figure 18:** Classifier type with Cases with T-L codes for AIS, Controls without T-L codes for AIS (TI) case-control combination varies in calibration success between stroke score and actual proportion of patients at each probability.

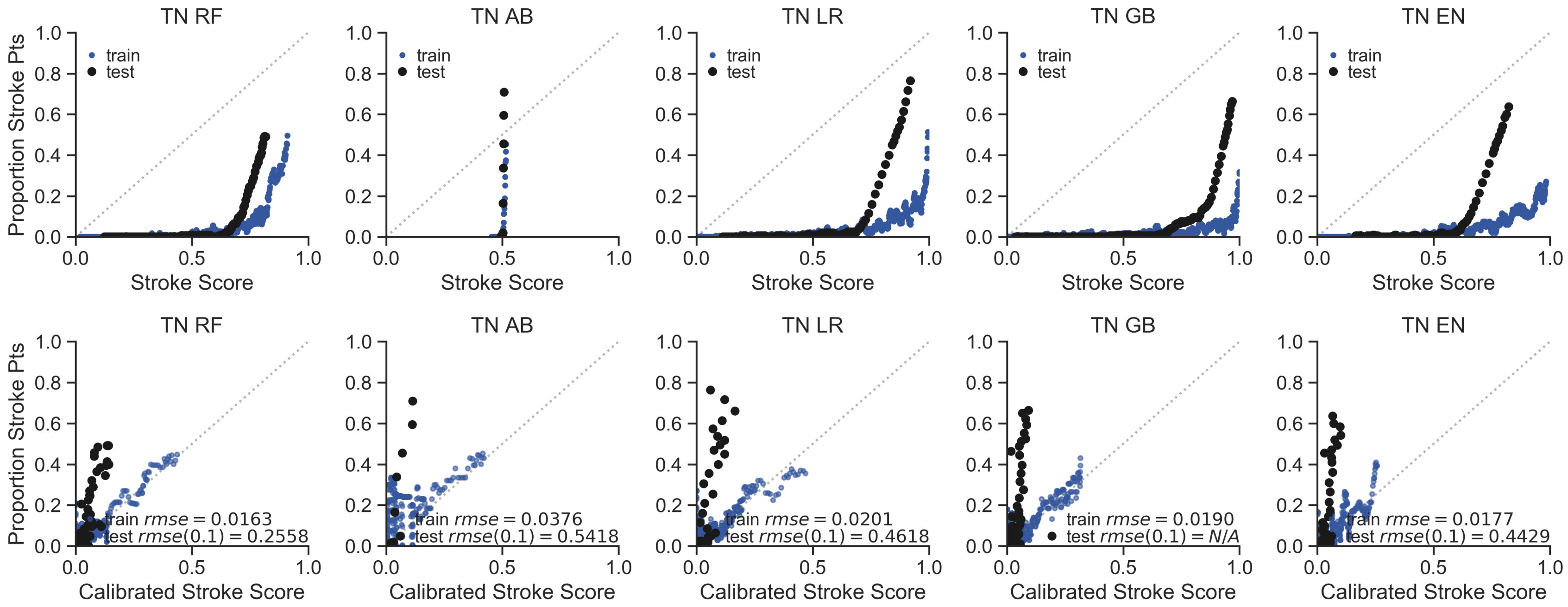


**Supplementary Figure 19:** Classifier type with Cases with T-L codes for AIS, Stroke Mimetic Controls (TN) case-control combination varies in calibration success between stroke score and actual proportion of patients at each probability.

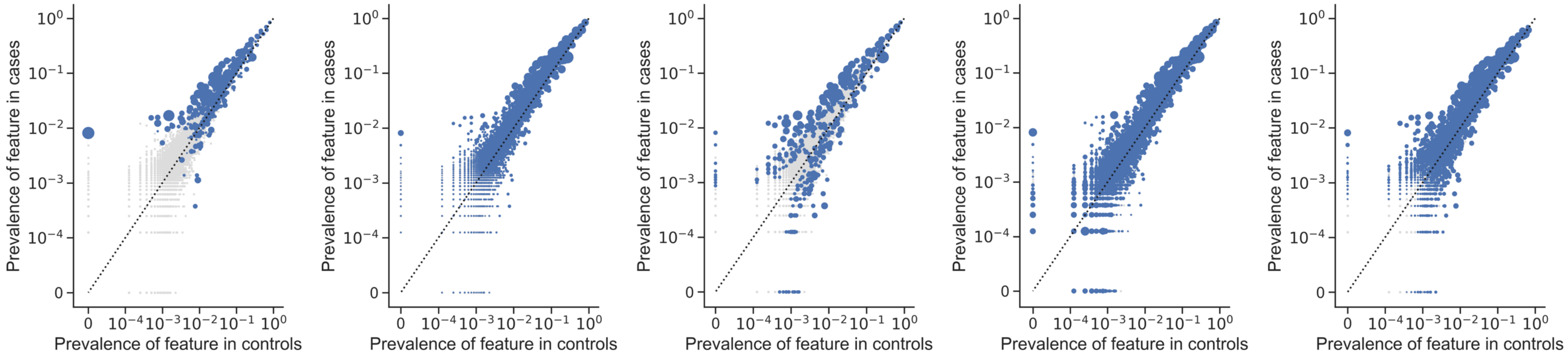


**Supplementary Figure 20:** Prevalence of features in cases vs controls in the TCI model. Left to Right: LR, RF, AB, GB, and EN models. Axes are on a logarithmic scale. Increasing size of blue dot correlates with higher feature importance or beta coefficient weight, depending on the classifier type. Gray dots are features with zero importance.

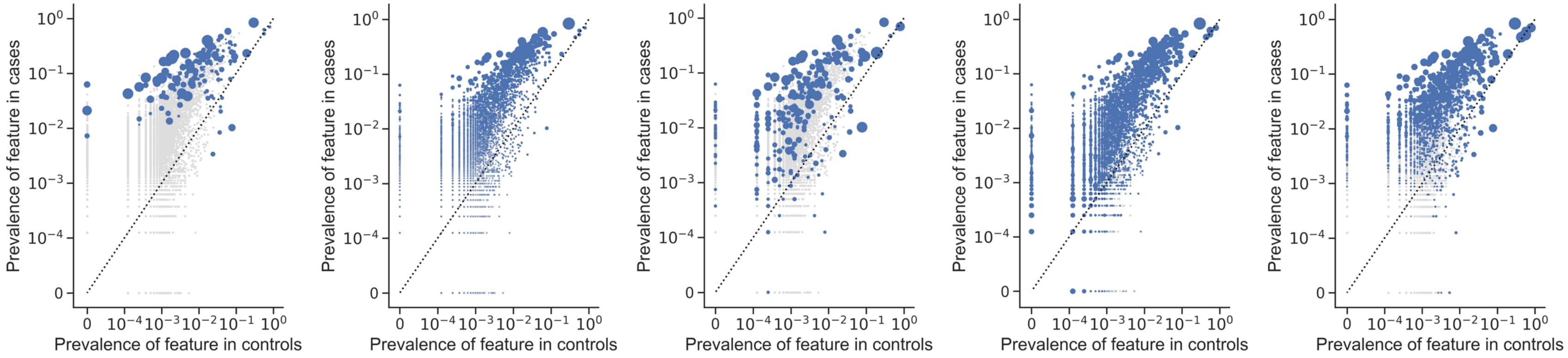


**Supplementary Figure 21:** Prevalence of features in cases vs controls in the TC model. Left to Right: LR, RF, AB, GB, and EN models. Axes are on a logarithmic scale. Increasing size of blue dot correlates with higher feature importance or beta coefficient weight, depending on the classifier type. Gray dots are features with zero importance.

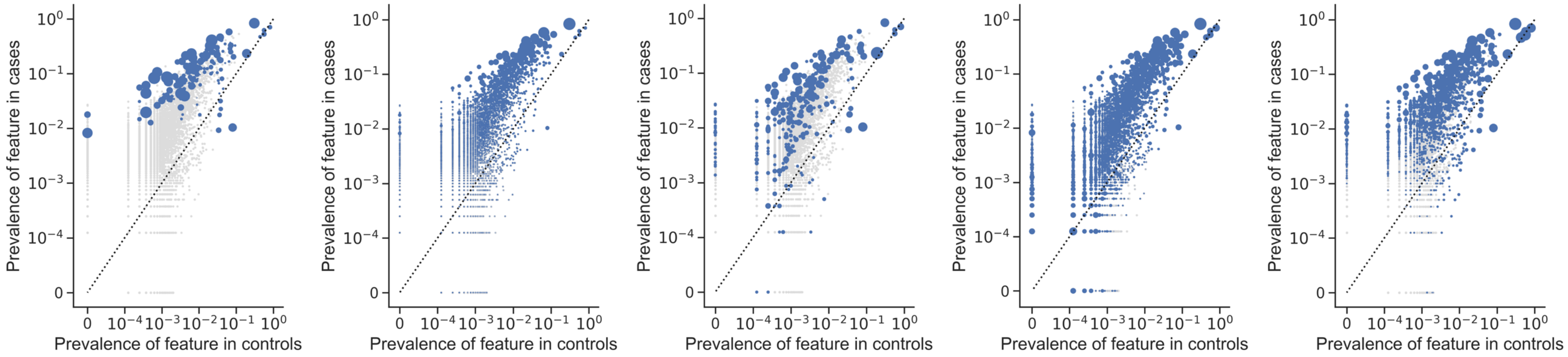


**Supplementary Figure 22:** Prevalence of features in cases vs controls in the TI model. Left to Right: LR, RF, AB, GB, and EN models. Axes are on a logarithmic scale. Increasing size of blue dot correlates with higher feature importance or beta coefficient weight, depending on the classifier type. Gray dots are features with zero importance.

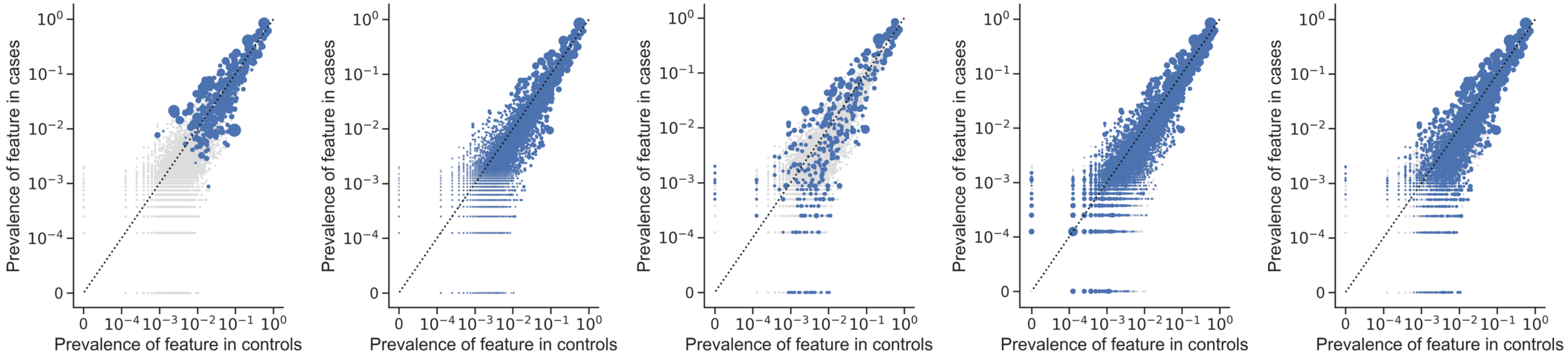


**Supplementary Figure 23:** Prevalence of features in cases vs controls in the TR model. Left to Right: LR, RF, AB, GB, and EN models. Axes are on a logarithmic scale. Increasing size of blue dot correlates with higher feature importance or beta coefficient weight, depending on the classifier type. Gray dots are features with zero importance.

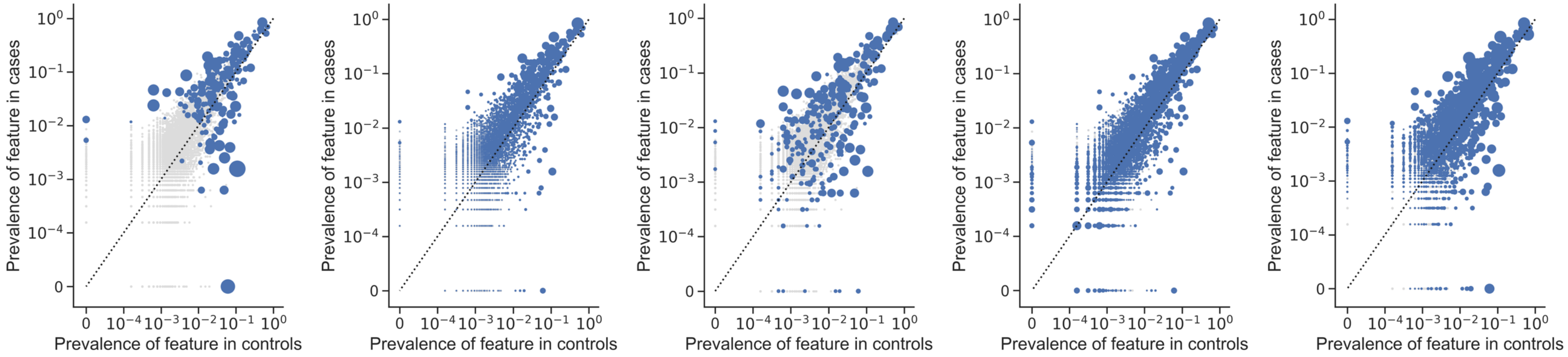


**Supplementary Figure 24:** Prevalence of features in cases vs controls in the TN model. Left to Right: LR, RF, AB, GB, and EN models. Axes are on a logarithmic scale. Increasing size of blue dot correlates with higher feature importance or beta coefficient weight, depending on the classifier type. Gray dots are features with zero importance.

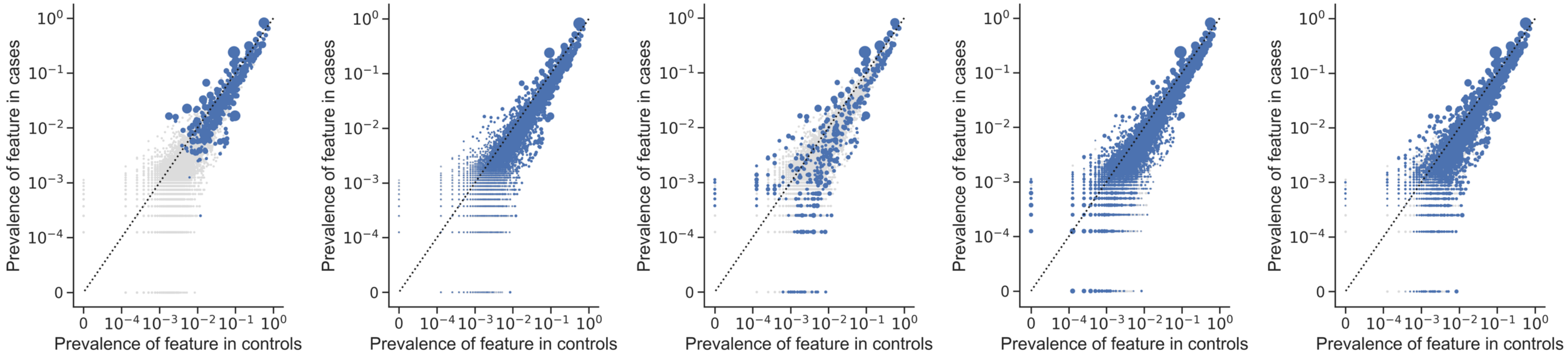


**Supplementary Figure 25:** Prevalence of features in cases vs controls in the CR model. Left to Right: LR, RF, AB, GB, and EN models. Axes are on a logarithmic scale. Increasing size of blue dot correlates with higher feature importance or beta coefficient weight, depending on the classifier type. Gray dots are features with zero importance.

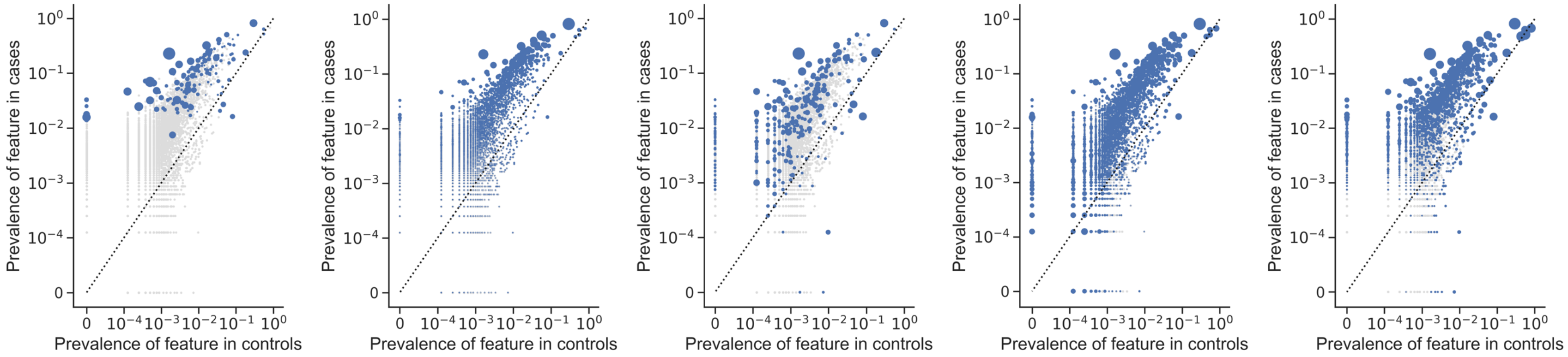


**Supplementary Figure 26:** Prevalence of features in cases vs controls in the CC model. Left to Right: LR, RF, AB, GB, and EN models. Axes are on a logarithmic scale. Increasing size of blue dot correlates with higher feature importance or beta coefficient weight, depending on the classifier type. Gray dots are features with zero importance.

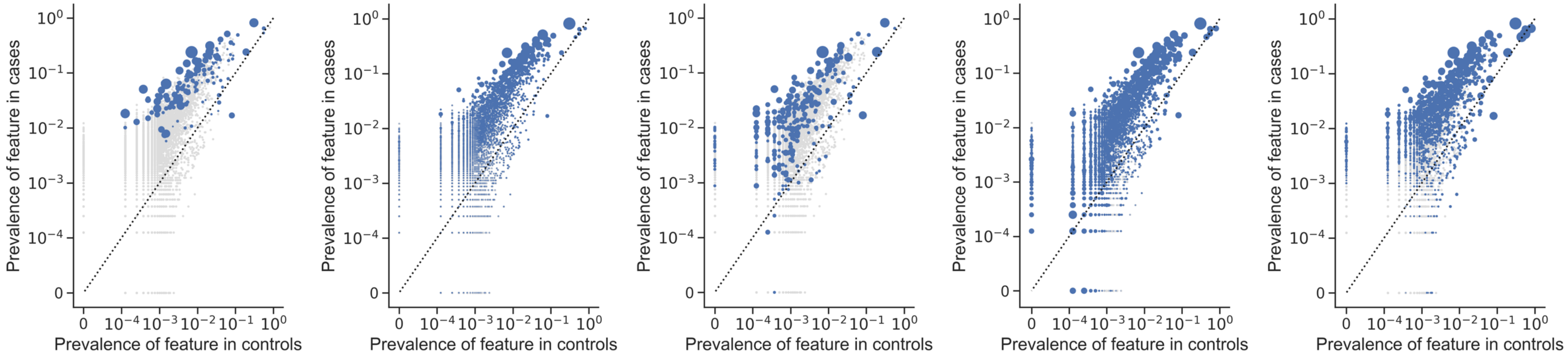


**Supplementary Figure 27:** Prevalence of features in cases vs controls in the CI model. Left to Right: LR, RF, AB, GB, and EN models. Axes are on a logarithmic scale. Increasing size of blue dot correlates with higher feature importance or beta coefficient weight, depending on the classifier type. Gray dots are features with zero importance.

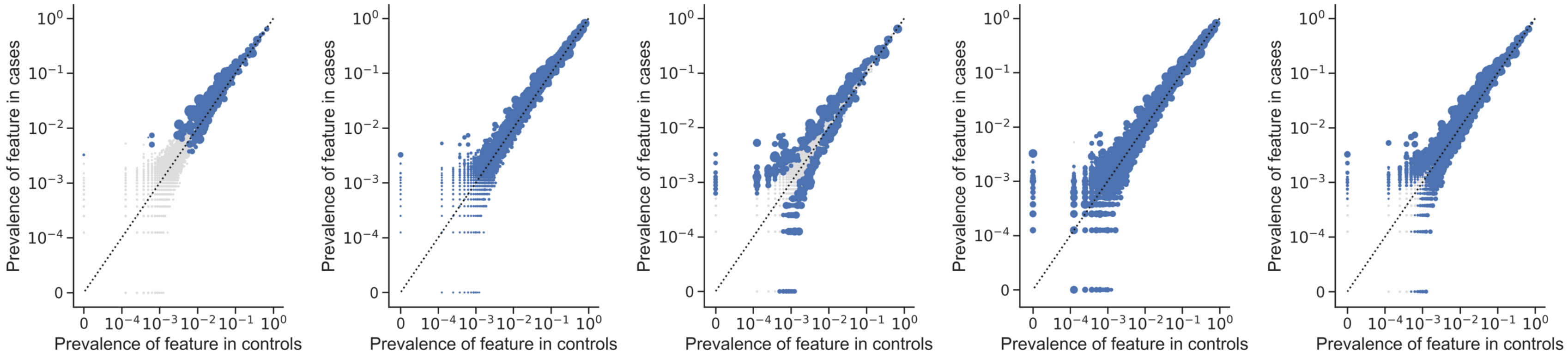


**Supplementary Figure 28:** Prevalence of features in cases vs controls in the CCI model. Left to Right: LR, RF, AB, GB, and EN models. Axes are on a logarithmic scale. Increasing size of blue dot correlates with higher feature importance or beta coefficient weight, depending on the classifier type. Gray dots are features with zero importance.

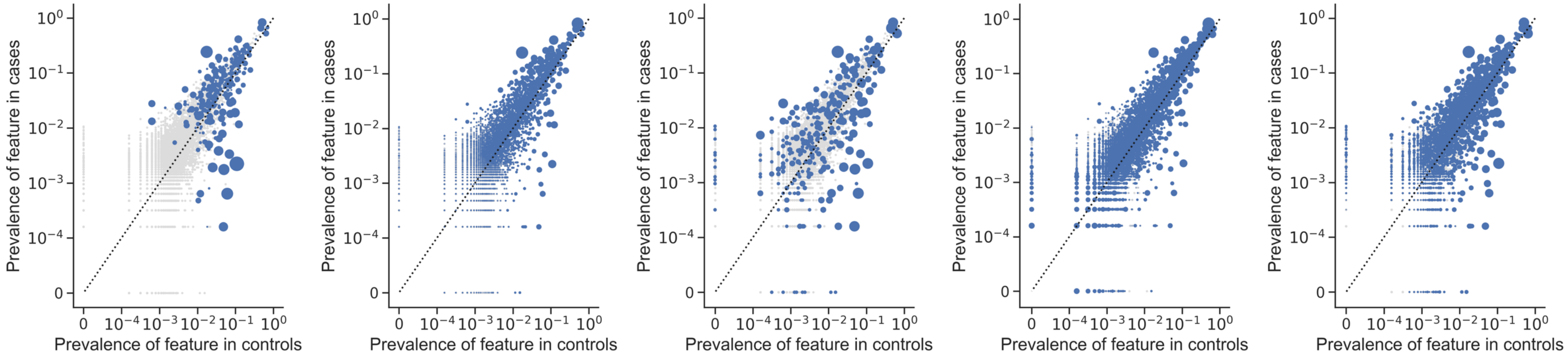


**Supplementary Figure 29:** Prevalence of features in cases vs controls in the CN model. Left to Right: LR, RF, AB, GB, and EN models. Axes are on a logarithmic scale. Increasing size of blue dot correlates with higher feature importance or beta coefficient weight, depending on the classifier type. Gray dots are features with zero importance.

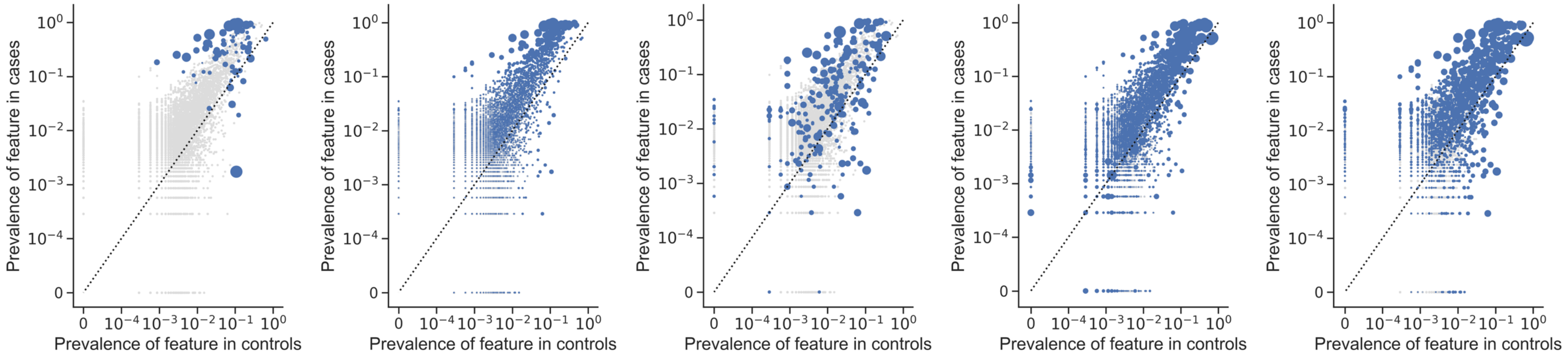


**Supplementary Figure 30:** Prevalence of features in cases vs controls in the SN model. Left to Right: LR, RF, AB, GB, and EN models. Axes are on a logarithmic scale. Increasing size of blue dot correlates with higher feature importance or beta coefficient weight, depending on the classifier type. Gray dots are features with zero importance.

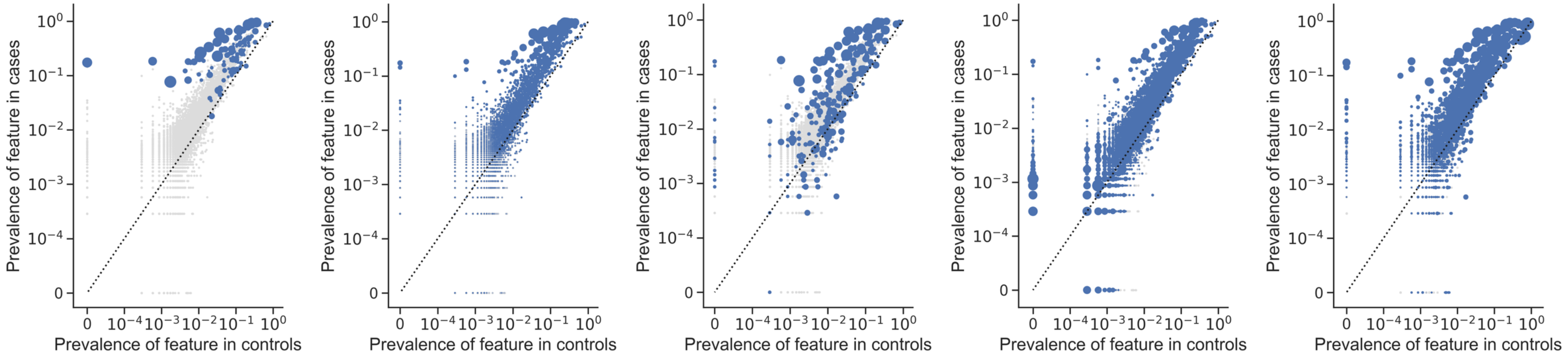


**Supplementary Figure 31:** Prevalence of features in cases vs controls in the SCI model. Left to Right: LR, RF, AB, GB, and EN models. Axes are on a logarithmic scale. Increasing size of blue dot correlates with higher feature importance or beta coefficient weight, depending on the classifier type. Gray dots are features with zero importance.

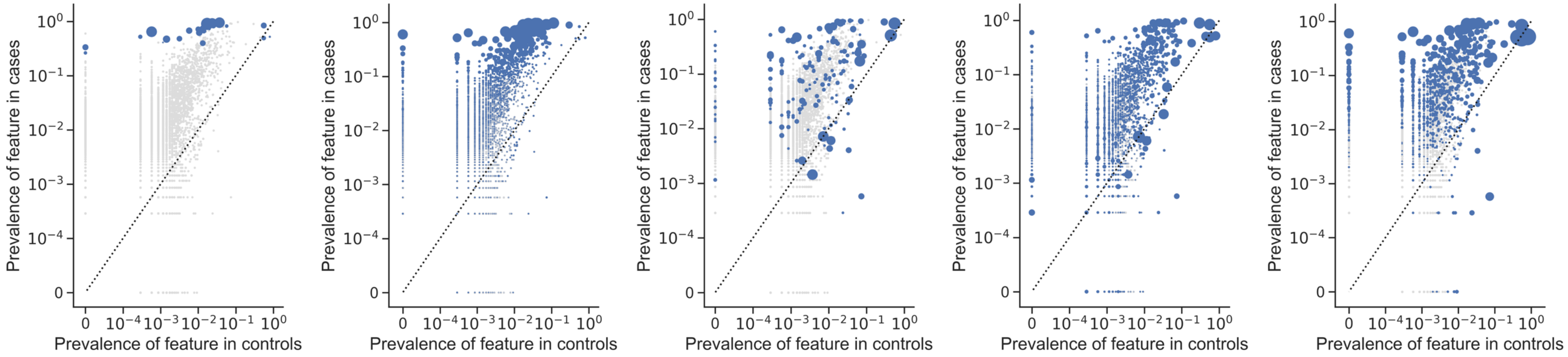


**Supplementary Figure 32:** Prevalence of features in cases vs controls in the SC model. Left to Right: LR, RF, AB, GB, and EN models. Axes are on a logarithmic scale. Increasing size of blue dot correlates with higher feature importance or beta coefficient weight, depending on the classifier type. Gray dots are features with zero importance.

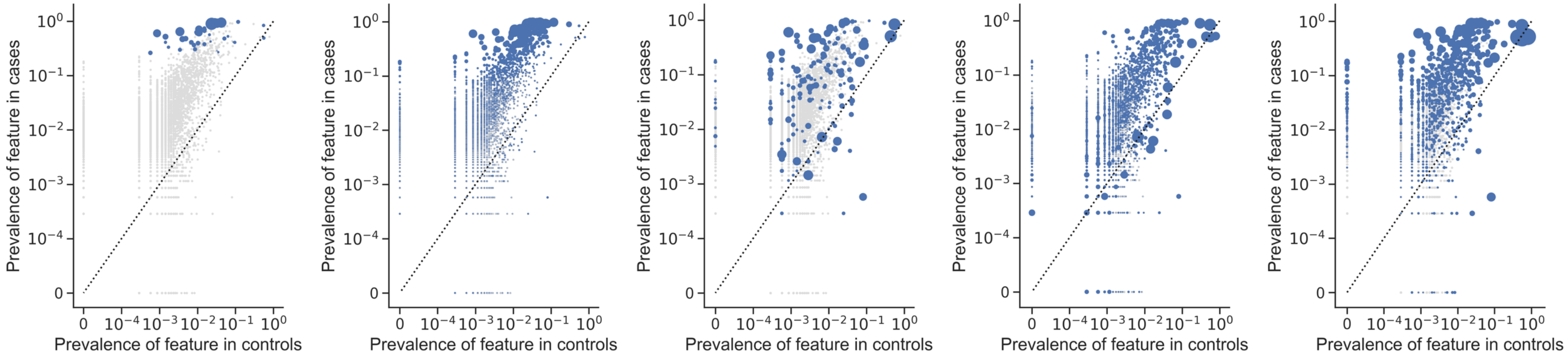


**Supplementary Figure 33:** Prevalence of features in cases vs controls in the SI model. Left to Right: LR, RF, AB, GB, and EN models. Axes are on a logarithmic scale. Increasing size of blue dot correlates with higher feature importance or beta coefficient weight, depending on the classifier type. Gray dots are features with zero importance.

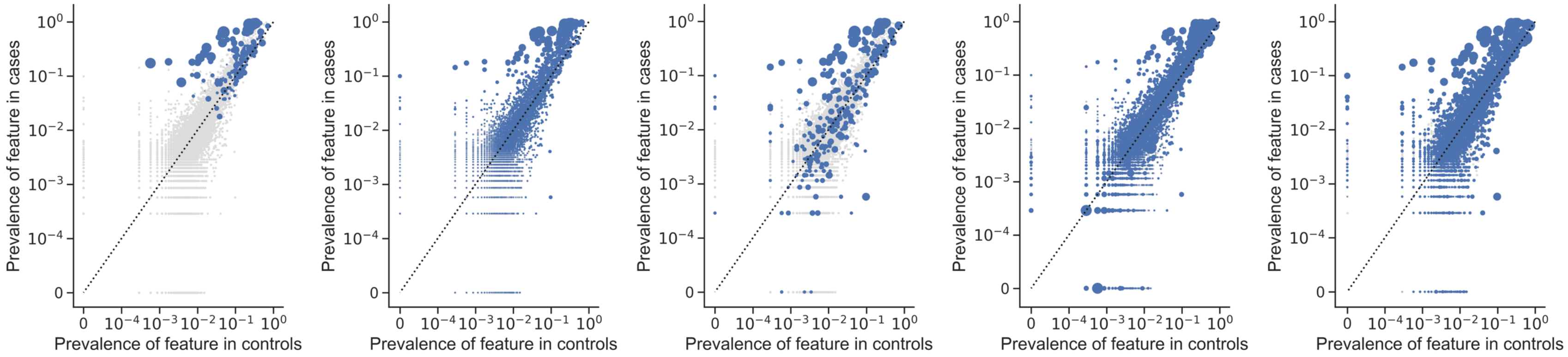


**Supplementary Figure 34:** Prevalence of features in cases vs controls in the SR model. Left to Right: LR, RF, AB, GB, and EN models. Axes are on a logarithmic scale. Increasing size of blue dot correlates with higher feature importance or beta coefficient weight, depending on the classifier type. Gray dots are features with zero importance.
